# Liebig’s law of the minimum in the TGF-β/SMAD pathway

**DOI:** 10.1101/2023.07.10.548398

**Authors:** Yuchao Li, Difan Deng, Chris Tina Höfer, Jihye Kim, Won Do Heo, Quanbin Xu, Xuedong Liu, Zhike Zi

**Affiliations:** Max Planck Institute for Molecular Genetics, Otto Warburg Laboratory, 14195 Berlin, Germany; German Federal Institute for Risk Assessment, Department of Experimental Toxicology and ZEBET, 10589 Berlin, Germany; Department of Biological Sciences, KAIST Institute for the BioCentury, Korea Advanced Institute of Science and Technology (KAIST), Daejeon, 305701 Republic of Korea; Department of Biochemistry, 3415 Colorado Ave, University of Colorado Boulder, Boulder, Colorado 80303-0596, United States; Key Laboratory of Quantitative Synthetic Biology, Shenzhen Institute of Synthetic Biology, Shenzhen Institute of Advanced Technology, Chinese Academy of Sciences, Shenzhen 518055, China

**Keywords:** systems biology, mathematical modeling, TGF-β pathway, cellular heterogeneity, synthetic biology

## Abstract

Cells use signaling pathways to sense and respond to their environments. The transforming growth factor-β (TGF-β) pathway produces context-specific responses. Here, we combined modeling and experimental analysis to study the dependence of the output of the TGF-β pathway on the abundance of signaling molecules in the pathway. We showed that the TGF-β pathway processes the variation of TGF-β receptor abundance using Liebig’s law of the minimum, meaning that the output-modifying factor is the signaling protein that is most limited, to determine signaling responses across cell types and in single cells. We found that the abundance of either the type I (TGFBR1) or type II (TGFBR2) TGF-β receptor determined the responses of cancer cell lines, such that the receptor with relatively low abundance dictates the response. Furthermore, nuclear SMAD2 signaling correlated with the abundance of TGF-β receptor in single cells depending on the relative expression levels of TGFBR1 and TGFBR2. A similar control principle could govern the heterogeneity of signaling responses in other signaling pathways.

**One-sentence summary:** Heterogeneous TGF-β signaling responses are dictated by the low abundance TGF-β receptor in different cell types and in single cells, resembling Liebig’s law of the minimum.

## Introduction

Cells sense and respond to their environment through signaling pathways. These pathways convert extracellular factors into signaling activities, resulting in the responses such as regulation of gene expression and determination of cell fate. Heterogeneous signaling responses to the same environmental cue often occur in different cell types and in individual cells. Intrinsic noise and extrinsic noise contribute to cell-to-cell variability (Eldar & Elowitz, 2010; Eling *et al*, 2019; Losick & Desplan, 2008; Mitchell & Hoffmann, 2018; Niepel *et al*, 2009; Snijder & Pelkmans, 2011). Intrinsic noise reflects the stochastic fluctuations of biochemical reactions, whereas extrinsic noise originates from differences in protein abundance, cell cycle stage, and other parameters. Cellular protein abundances have a large dynamic range with several orders of magnitude (Buccitelli & Selbach, 2020). Variation in protein abundance affects the heterogeneity of signaling responses (Cohen-Saidon *et al*, 2009; Filippi *et al*, 2016; Gabor *et al*, 2021; Kramer *et al*, 2022; Lun *et al*, 2017; Strasen *et al*, 2018; Wu *et al*, 2021).

The transforming growth factor-β (TGF-β) pathway regulates many fundamental processes, including cell proliferation, differentiation, and migration (Shi & Massague, 2003). The effects of TGF-β depend on the cellular context (Massague, 2012; Zhang, 2018), providing a useful system to investigate the relationship between signaling protein abundance and signaling output. The core of the TGF-β pathway is composed of TGF-β receptors (TGFBR1, TGFBR2) that perceive the ligand TGF-β, receptor-regulated SMADs (R-SMADs: SMAD2, SMAD3) and common SMAD (Co-SMAD: SMAD4) effectors that mediate the transcriptional response. Additionally, inhibitory SMAD (I-SMAD: SMAD7) plays a crucial role in negatively regulating this pathway. The intricacies of the TGF-β pathway extend further with complex feedback regulations and interactions with various other signaling pathways (Derynck & Budi, 2019; Luo, 2017; Zhang, 2017).

At receptor level, TGF-β receptors undergo tight regulation through processes such as endocytosis and posttranslational modifications (Budi *et al*, 2016). The activities of SMAD proteins, undergoing nucleocytoplasmic shuttling, are controlled by upstream ligand-receptor complexes, the phosphorylation events at the linker region, as well as actions of SMAD phosphatases (Bruce & Sapkota, 2012; Gao *et al*, 2009; Hill, 2009; Kamato & Little, 2020; Lin *et al*, 2016). This complexity contributes to the enigmatic dual roles played by TGF-β in cancer progression. While functioning as a tumor suppressor in the early stages, TGF-β can promote tumor progression at later stages. Changes in the expression of TGF-β receptors or downstream signaling molecules can switch its role from tumor suppressor to tumor promoter. Therefore, owing to its diverse effects in different stages of cancer, targeting TGF-β for therapeutic intervention poses a substantial challenge.

Previous computational modeling studies have contributed to our understanding of how SMAD signaling is regulated by factors such as the timing and doses of the TGF-β ligand, receptor dynamics, feedback regulations, SMAD shuttling and oligomerization (Lucarelli *et al*, 2018; Schmierer *et al*, 2008; Sorre *et al*, 2014; Vilar *et al*, 2006; Vizan *et al*, 2013; Zi *et al*, 2011; Zi & Klipp, 2007). The abundance of the core components and feedback regulators could determine TGF-β responses by regulating SMAD signaling in the nucleus (Derynck & Budi, 2019; Massague, 2012). Heterogeneous SMAD signaling responses in single cells can arise from the variability in protein levels (Frick *et al*, 2017; Strasen *et al*., 2018). For the bone morphogenetic protein (BMP) pathway, diverse response profiles depend on the combinations of BMP ligands with different receptor binding affinities (Antebi *et al*, 2017; Klumpe *et al*, 2022; Mattiazzi Usaj *et al*, 2021). Manipulation of the abundance of BMP receptor subunits reprograms response profiles for different BMP ligand pairs (Antebi *et al*., 2017). The same TGF-β ligand can initiate different signaling responses depending on the cellular context, but the underlying control principle remains unclear.

In this work, we examined how differences in signaling protein abundance dictates TGF-β signaling response profiles in different cell lines and in single cells using a combination of computational and experimental approaches. Computational modeling revealed that SMAD signaling responses can be determined by the abundance of the type I or type II TGF-β receptor. The type of TGF-β receptor present in relatively low numbers played a dominant role in shaping signaling responses, which resembles Liebig’s law of the minimum (Liebig, 1840). Further experiments showed that different cell types have distinct response sensitivities to the variation of TGF-β receptors, depending on the relative expressions of TGFBR1 and TGFBR2. In addition, we found that the nuclear SMAD2 signaling strongly correlates with the expression of either of the TGF-β receptors in single cells, depending on the expression pattern of the receptors. Together, these results revealed an effect of the minimum control in the TGF-β pathway, which may be an important principle of control in signaling pathways with context-dependent outputs.

## Results

### TGF-β receptors display heterogeneous expression profiles across diverse cell types

Utilizing the Human Protein Atlas data (Karlsson *et al*, 2021), we first examined the transcript expression profiles of TGF-β receptors across various cell types. Among 1055 cell lines, both type I (*TGFBR1*) and type II (*TGFBR2*) TGF-β receptors displayed large variations (Figure 1A). The *TGFBR1*-to-*TGFBR2* expression ratios were also heterogeneous in these cell lines (Figure 1B). Numerous cancer cell lines had imbalanced expression levels of *TGFBR1* and *TGFBR2*. For instance, RH-30, TE-4 and NCI-H524 considerably higher levels of *TGFBR1* compared to *TGFBR2*, while HepG2, RT-4, and KP-4 had elevated *TGFBR2* expression relative to *TGFBR1*. Importantly, previous work has demonstrated a high correlation between protein and RNA data for TGFBR1 and TGFBR2 within the same cell line (Spearman’s correlation: 0.672 for TGFBR1, 0.771 for TGFBR2), based on comprehensive data from 375 cancer cell lines (Nusinow *et al*, 2020). This robust correlation supports the likelihood of imbalanced protein expression of TGFBR1 and TGFBR2 in cancer cell lines.

**Figure 1:**
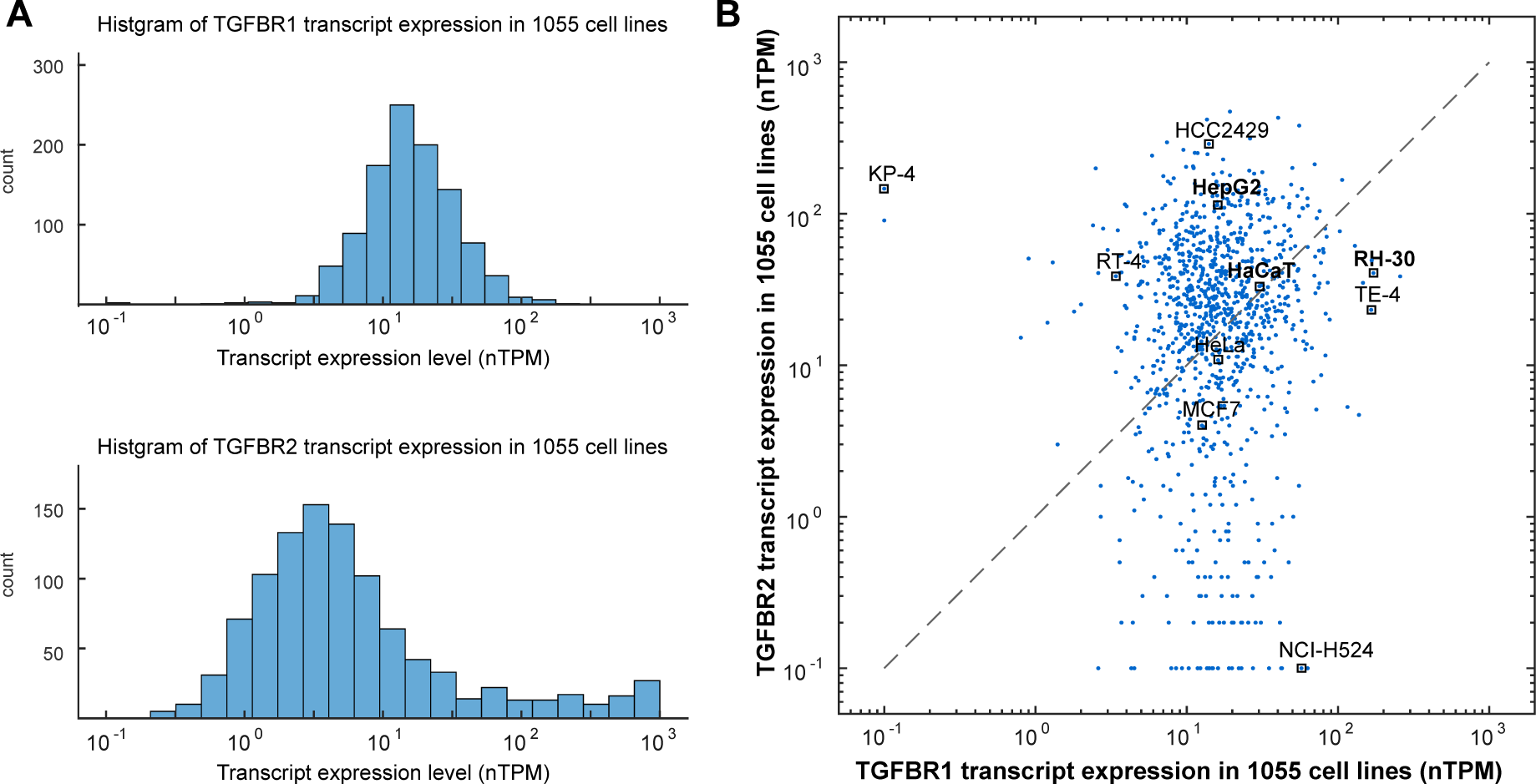
Transcript Expression Profile of TGFBR1 and TGFBR2 in 1055 Cell Lines. The transcript data for TGFBR1 and TGFBR2 were obtained from the Human Protein Atlas database version 22.0. The expression levels are represented in normalized transcripts per million (nTPM). (A) Histogram depicting the distribution of TGF-β receptors expression across various cell lines. (B) Imbalanced expression of TGFBR1 and TGFBR2 was observed in numerous cancer cell lines. Each dot in the plot represents the expression levels of TGFBR1 and TGFBR2 in one of the 1055 cell lines. Specific cell lines of interest are labeled as representatives.

### A minimal model predicts Liebig’s law of the minimum in the TGF-β pathway

The heterogeneous expression profile of TGF-β receptors inspired us to analyze how different types of cells to process the variations of TGF-β receptors. To address this question, we first developed a minimal mathematical model that describes the key aspects of the TGF-β pathway with two reactions (Figure 2A, Supplementary Information). The first reaction represents the binding of the TGF-β ligand (*L*) to type I (*R*1) and type II (*R*2) TGF-β receptors, which forms the ligand-receptor complex (*LRC*). The second reaction describes the phosphorylation and dephosphorylation of the SMAD protein. As seen in the mathematical models of the BMP pathway (Antebi *et al*., 2017; Su *et al*, 2022), the minimal model assumes that the phosphorylation of the SMAD protein is proportional to the activated ligand-receptor complex and the total amount of the species in the model remains constant (conservation of mass).

**Figure 2:**
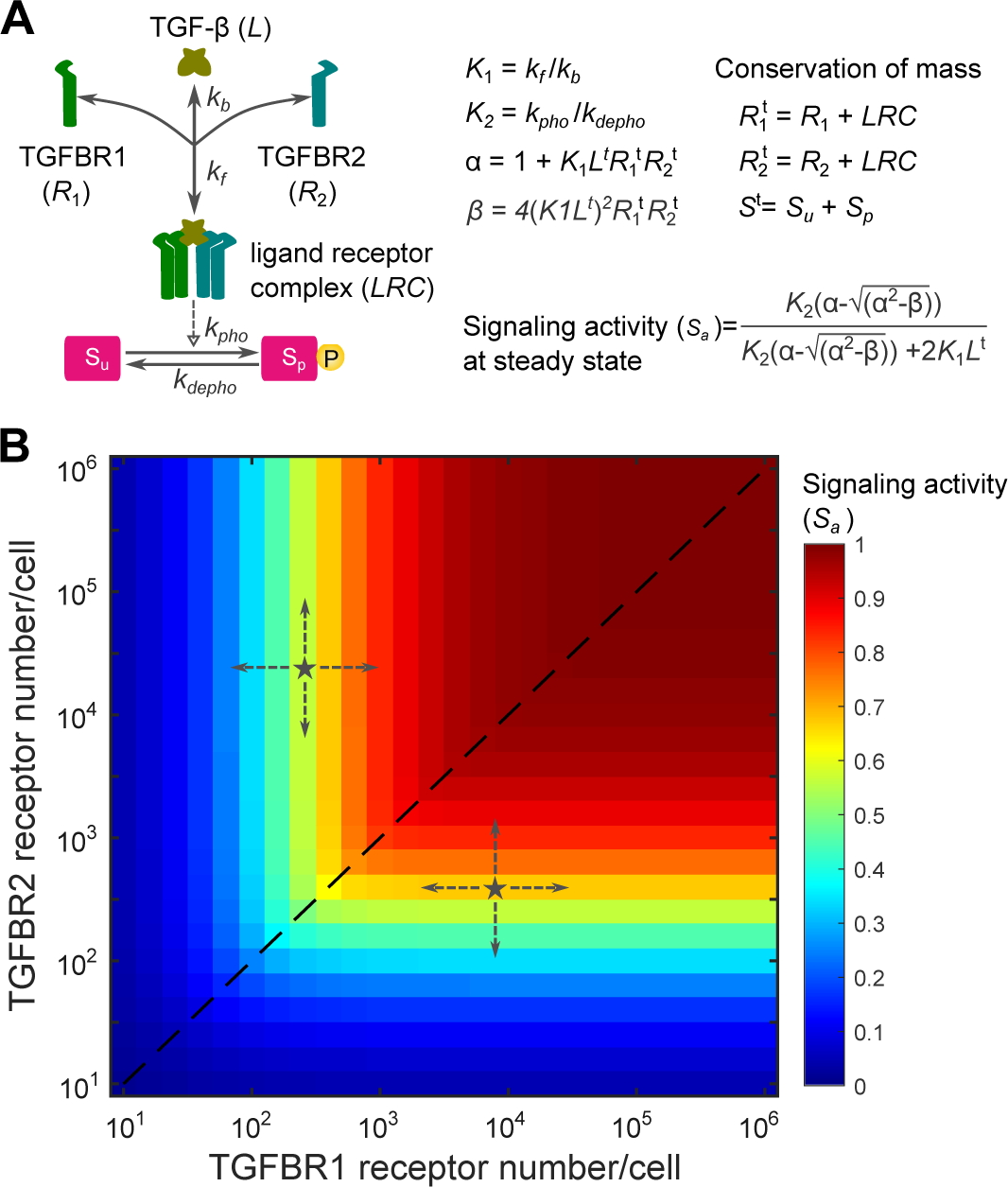
A minimal mathematical model predicts that TGF-β signaling activity is dictated by the type of TGF-β receptor in lower abundance. (A) A minimal model of the TGF-β pathway with two reactions including the TGF-β ligand (*L*), which interact with type I receptor (*R*1) and type II receptor (*R*2) with a binding affinity (*K*1). The active ligand-receptor complex (*LRC*) phosphorylates the SMAD protein with an equilibrium coefficient (*K*2). TGF-β signaling activity (*S*a) at the steady state can be analytically calculated with equations (refer to Supplementary Information for model details and equation derivation). (B) The response profile of TGF-β signaling activity (*S*a) at the steady state is shown in relation to a 2-dimensional matrix of TGFBR1 and TGFBR2 expression numbers spanning five orders of magnitude. *K*1=1000, *K*2=10. TGF-β ligand concentration is set to be 0.1 nM.

Using this simple model, we investigated the impact of TGF-β receptor variations on steady-state TGF-β signaling activity (*Sa*), which has an analytic solution (Supplementary Information). To quantitively characterize the signal response, we analyzed the signaling activity (*Sa*) with respect to a 2-dimensional matrix of TGFBR1 and TGFBR2 numbers spanning 5 magnitudes. The TGF-β signaling activity profile revealed a notable pattern: signaling activity was primarily dictated by the receptor with lower abundance (Figure 2B). When TGFBR1 and TGFBR2 were equally expressed, TGF-β signaling activity was proportional to the receptor numbers (the dashed line in Figure 2B). Interestingly, when the amounts of TGFBR1 and TGFBR2 were imbalanced (denoted as stars in Figure 2B), TGF-β signaling is sensitive to changes of the TGF-β receptor with lower abundance, while remaining robust to changes of the TGF-β receptor with higher abundance. This observation aligns with Liebig’s law of the minimum, which states that the growth rate of a species is determined by the scarcest resource (Liebig, 1840). It is noteworthy that similar response profiles were predicted when setting a high binding affinity (*K1*) for ligand-receptor interactions in the minimal model (Figure S1A). However, with a small binding affinity, the minimal model indicates that signal response remains proportional to the product of TGFBR1 and TGFBR2 abundance (Figure S1B). This suggests that the observed response patterns aligning with Liebig’s law of the minimum depend on binding affinity of ligand-receptor interactions in the minimal model.

### An extended model further demonstrates that TGF-β signaling is governed by the type of TGF-β receptor in lower abundance

For simplicity, the above-mentioned minimal model made several assumptions by neglecting specific features of the TGF-β pathway such as receptor internalization and recycling, nuclear import and export of SMAD protein, and the feedback regulations. Consequently, it remains unclear whether the predicted TGF-β signaling response profiles aligning with Liebig’s law of the minimum holds true in a more intricate TGF-β pathway when accounting for additional signaling processes and regulations. To address this uncertainty, we developed an extended mathematical model that characterizes TGF-β signaling processes with greater complexity, including ligand-receptor interaction, receptor endocytosis, SMAD2 nucleocytoplasmic shuttling, and negative feedback regulation (Figure 3A). For simplicity, non-SMAD signaling and crosstalk between TGF-β/SMAD and other signaling pathways were not considered in this model. Building upon our previous model (Zi *et al*., 2011), the extended model employs mass action-based ordinary differential equations to describe the dynamics of the TGF-β pathway (Supplementary Information).

**Figure 3.**
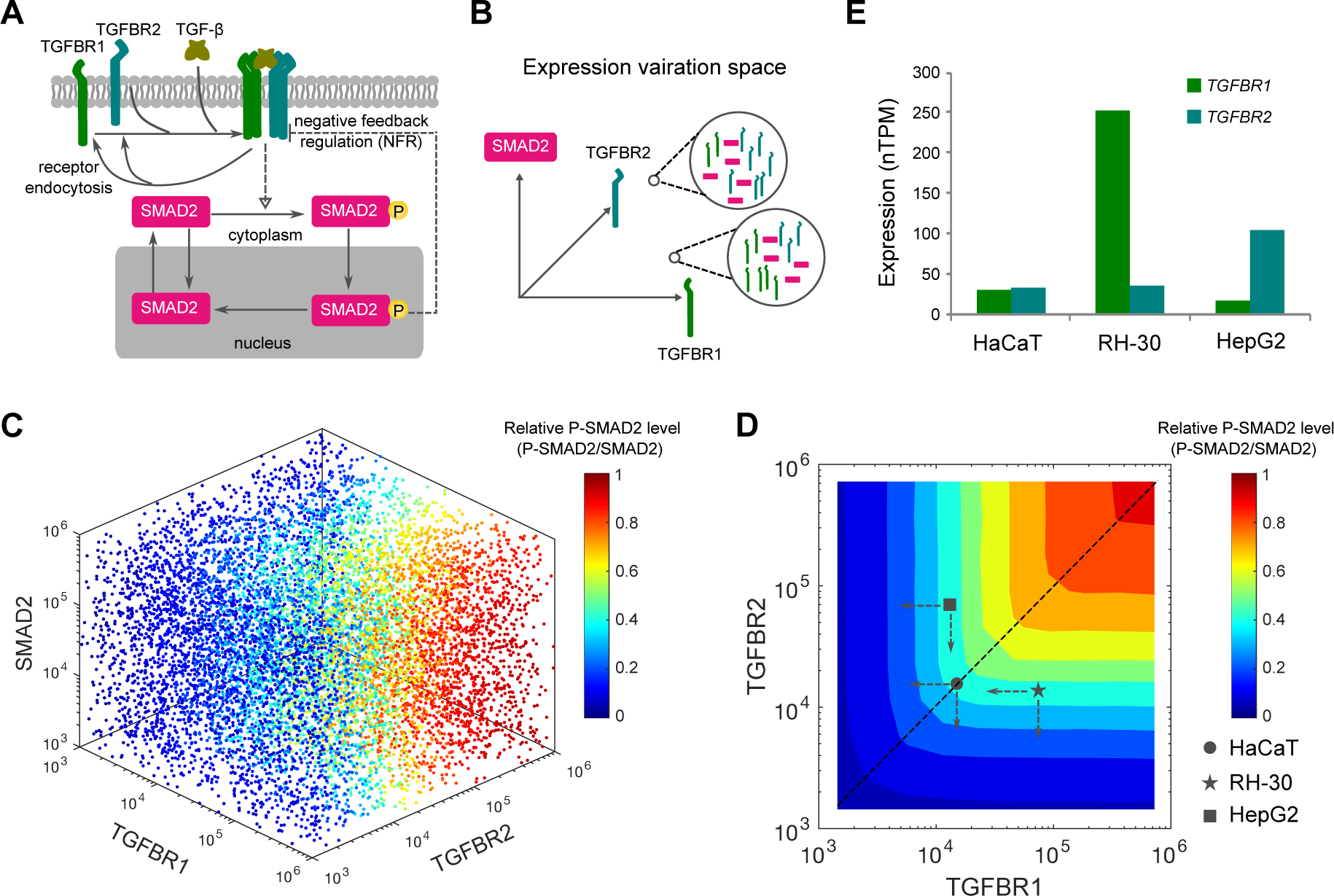
Computational modeling of heterogeneous signaling responses in the variation space of TGF-β receptors and SMAD2 expression. (A) Scheme of the extended TGF-β signaling model. The development of the extended model is described in the Supplementary Information. (B) Heterogeneous expression of TGF-β receptors and SMAD2 is illustrated as points in 3 multi-dimensional spaces. The two magnified circles exemplify two possible expression profiles of TGFBR1, TGFBR2 and SMAD2. (C) Model predictions for normalized P-SMAD2 responses in the expression space of TGFBR1, TGFBR2 and SMAD2. Each scatter point indicates a random set of protein expression levels in the space. The point colors indicate the normalized P-SMAD2 responses (P-SMAD2/SMAD2). (D) Contour landscape of the normalized P-SMAD2 responses to the combinations of TGFBR1 and TGFBR2. The dashed line indicates relationship between the relative P-SMAD2 level and the receptors when equal amounts of each receptor are present. The abundances of TGFBR1 and TGFBR2 in HaCaT, RH-30 and HepG2 cell lines are mapped according to the relative expression data shown in Figure 3E and the measured HaCaT TGF-β receptor number shown in Figure S2B. The dashed arrows indicate the directions of receptor changes. (E) TGF-β receptor expression profiles for HaCaT, RH-30 and HepG2 cell lines. Bars indicate expression levels of each receptor (normalized transcripts per million, nTPM). The data is based on the Human Protein Atlas database version 22.0.

To parameterize the extended model, we conducted quantitative immunoblotting experiments using HaCaT cells. These experiments allowed us to measure the absolute abundance of SMAD2 and TGFBR2 (Figure S2), the rate of TGFBR2 internalization (Figure S3), and the half-life of SMAD2, TGFBR1, and TGFBR2. Using these experimental data, we determined the initial conditions and some model parameter values (Supplementary Information). To constrain the remaining nine unknown parameters, we fitted the model to 102 data points obtained from HaCaT cells exposed to varying concentrations of TGF-β. Notably, the extended model simulations accurately reproduced the experimental data, encompassing the dynamics of TGF-β in the medium, overall SMAD2 phosphorylation (P-SMAD2), cytoplasmic SMAD2 and P-SMAD2, as well as nuclear SMAD2 and P-SMAD2 (Figure S4). Profile likelihood analysis confirmed the practical identifiability of the estimated model parameters (Figure S5). Consequently, the extended model was rigorously refined with meticulous parameterization.

We next conducted simulations on the extended model to investigate the variations in signaling components among different types of cells. Specifically, we simultaneously varied the abundances of TGF-β receptors and SMAD2 in the extended model by randomly sampling from multi-dimensional spaces (Figure 3B). By repeating the model simulations and analyzing the SMAD2 signaling output for 100,000 variations of TGF-β receptors and SMAD2, we observed interesting findings. Our results showed that the abundance of TGF-β receptors, rather than SMAD2, played a predominant role in controlling the variations of relative P-SMAD2 (the amount of P-SMAD2 normalized to SMAD2 abundance) and normalized nuclear SMAD2 signaling (SMAD2 N2C fold change) (Figure 3C, Figure S6A).

To gain further insights into the specific patterns of SMAD signaling response resulting from different TGF-β receptor abundances, we plotted the two SMAD2 signaling outputs in relation to the abundance of TGFBR1 and TGFBR2 (Figure 3D, Figure S6B). Remarkably, the extended model predicted similar response patterns for relative P-SMAD2 and normalized nuclear SMAD signaling, reinforcing the idea that the TGF-β receptor with lower abundance acts as a key determinant of SMAD signaling response. Additional model simulations suggested that the low abundance of a TGF-β receptor acts as a bottleneck for SMAD2 signaling, with TGF-β stimulations at lower concentrations (Figure S7). Taken together, our findings from both the simplified and extended models support the notion that cells utilize a minority rule to process the variations in TGFBR1 and TGFBR2, ultimately influencing SMAD signaling output.

### TGF-β signaling is sensitive to perturbations of TGF-β receptor in lower abundance, but is robust to changes of TGF-β receptor in higher abundance

To experimentally test how SMAD signaling is changed with respect to the perturbations of TGF-β receptors in different cell types, we selected three cell lines with different ratios of TGFBR1 and TGFBR2: HaCaT cells with similar amounts of *TGFBR1* and *TGFBR2*, RH-30 cells with more *TGFBR1* than *TGFBR2*, and HepG2 cells with more *TGFBR2* than *TGFBR1* (Figure 3E). Notably, consistent imbalances in TGFBR1-to-TGFBR2 receptor ratios were observed in both HepG2 and RH30 cell lines, as corroborated by diverse sources of databases (Table S1). The relative abundances of TGFBR1 and TGFBR2 in these cell lines were confirmed by immunoblotting experiments (Figure S8).

If Liebig’s law of the minimum holds for the TGF-β pathway, our model simulations suggested that HaCaT, RH-30, and HepG2 cells will display distinct sensitivities in response to variations in TGFBR1 and TGFBR2 (Figure 3D). Notably, RH-30 cells demonstrate significantly lower expression levels of TGFBR2 compared to TGFBR1. Consequently, the computational model simulations predicted that SMAD2 signaling in RH-30 cells would be sensitive to alterations in TGFBR2, but not TGFBR1. On the other hand, HepG2 cells display a higher abundance of TGFBR2 relative to TGFBR1, leading the model to predict that SMAD2 signaling in these cells would be sensitive to the reduction of TGFBR1, rather than TGFBR2. Finally, given that HaCaT cells express comparable quantities of both receptors, the model predicts that SMAD2 signaling in these cells would exhibit sensitivity to the reduction of either receptor.

To qualitatively test these predictions, we compared the profiles of SMAD2 signaling to changes of TGFBR1 and TGFBR2 abundance in HaCaT, RH-30, and HepG2 cells. Using different concentrations of small interfering RNA (siRNA), we knocked down individual TGF-β receptors to varying amounts in these cell lines. Our data indicated that both TGFBR1 and TGFBR2 were effectively knocked down with the increasing concentrations of siRNAs of TGF-β receptors (Figure S9). Previous studies showed that SMAD2 phosphorylation peaked around 1 hour after addition of TGF-β to the media (Strasen *et al*., 2018; Zi *et al*., 2011). Therefore, we measured SMAD2 phosphorylation and TGF-β receptor amounts in the siRNA knockdown samples relative to the non-targeting control (ntc) siRNA samples at 1 hour after TGF-β stimulation (Figure 4, Figure S10). The experimental data validated each prediction of the model: in RH-30 cells, SMAD2 signaling is sensitive to the reduction of TGFBR2, but not TGFBR1. Conversely, in HepG2 cells, it displayed a notable dependence on the reduction of TGFBR1, but not TGFBR2. Lastly, in HaCaT cells, SMAD2 signaling demonstrated comparable sensitivity to knockdown of either receptor.

**Figure 4.**
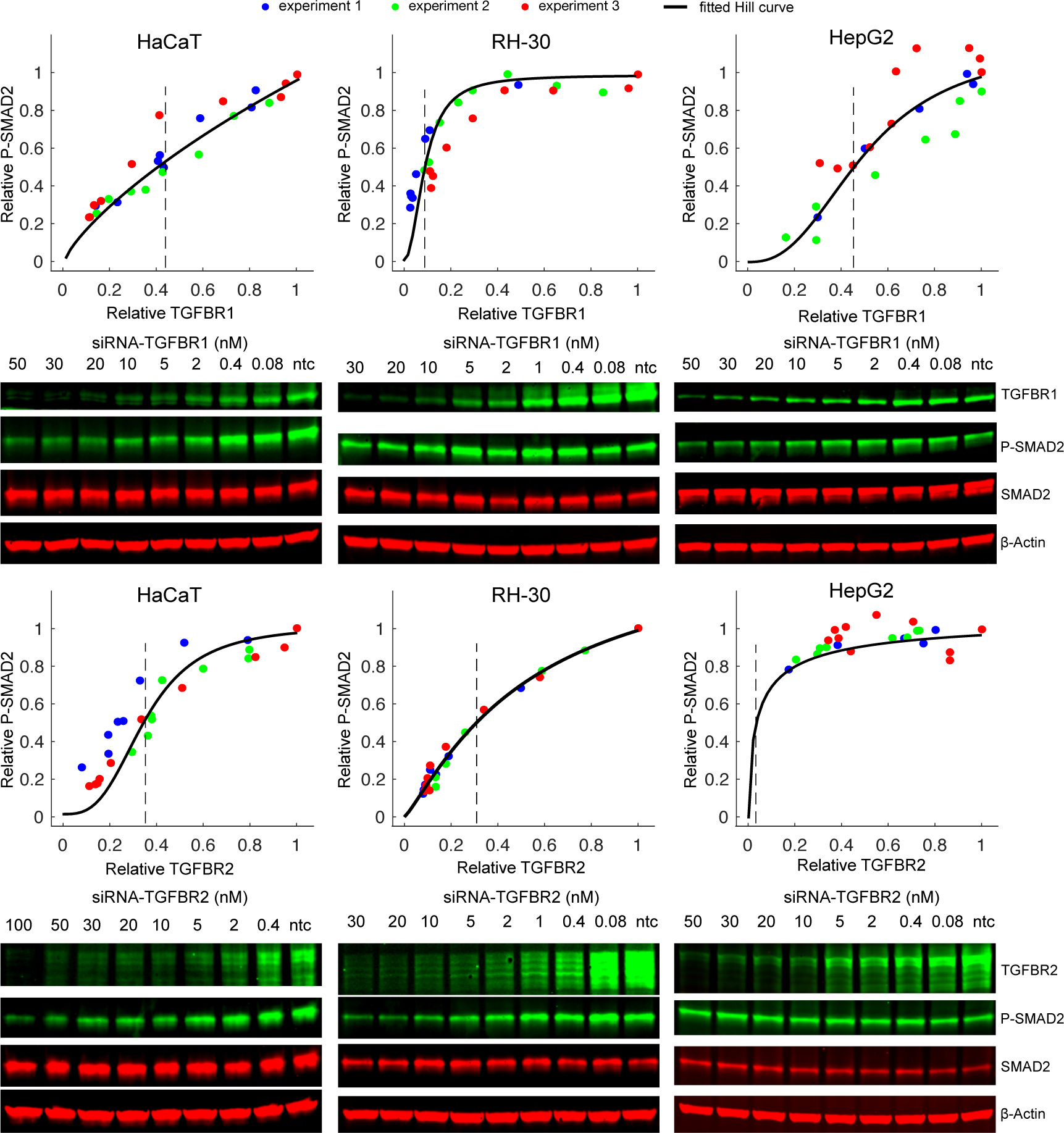
Different types of cells show distinct sensitivities to the knockdown of TGF-β receptors. Each type of TGF-β receptors was knocked down to different levels using various concentrations of siRNA. The relative P-SMAD2 responses and TGF-β receptor expression levels at 1 h after 100 pM TGF-β stimulation were measured and normalized to the non-targeting control (ntc) siRNA sample in the same experiment. Each plot contains the quantified data from three biological replicates. Immunoblotting results from one representative experiment are shown below each plot. The expression levels of relative TGFBR1 and TGFBR2 are plotted as a function of siRNA concentration in Figure S9. The black curve represents the fitted Hill curve and the dashed lines indicate the mean fold change of TGF-β receptor leads to the half of the P-SMAD2 response in the ntc sample (EC50). A statistical analysis of the fitted EC50s from 3 experiments is shown in Figure S10.

We then investigated whether TGF-β-inducible gene expression responses demonstrate similar sensitivities to the variations in the abundance of low-expressed TGF-β receptors in different cell lines. We knocked down TGF-β receptors using different concentrations of siRNAs and measured mRNA levels of TGF-β target genes (*SMAD7*, *PAI1* and *JUNB*) 1 h after TGF-β stimulation in HaCaT, RH30 and HepG2 cells. As shown in Figure S11, the expression of *SMAD7*, *PAI1* and *JUNB* were sensitive to the downregulation of TGFBR2 but not TGFBR1 in RH30 cells. In contrast, as TGFBR2 is more abundant than TGFBR1 in HepG2 cells, the expression of these genes was more sensitive to variations in TGFBR1 than to the changes in TGFBR2 levels. Since HaCaT cells express similar levels of TGFBR1 and TGFBR2, the expression of *SMAD7*, *PAI1* and *JUNB* showed similar sensitivities to the reduction of both receptors.

Overall, our results suggest that the type of TGF-β receptor with lower abundance acts as a limiting factor for TGF-β signaling. Furthermore, the data indicate that manipulation of the lower abundance TGF-β receptor might be an effective way to influence TGF-β signaling in different cellular contexts.

### SMAD2 signaling correlates with the low abundance TGF-β receptor in single cells

Because individual cells can have different amounts of TGF-β signaling proteins, we explored the TGF-β signaling responses in single cells by performing single-cell model simulations (Figure 5A). The single-cell models had the same molecular interactions and kinetic parameter values as the population model, but signaling protein concentrations (TGFBR1, TGFBR2, SMAD2) were varied according to log-normal distributions. The variations of TGF-β receptors were achieved by setting the production rate as a constant with a value randomly generated from log-normal distributions. The mean value of each log-normal distribution is the same as the corresponding value in the population model. To mimic the variation of the strength of negative feedback regulation (NFR), the parameter (k_NFR) for the constant representing ligand-induced receptor degradation was sampled from a log-normal distribution. A coefficient of variation (CV) of 0.1 was applied for the variations of signaling proteins and NFR strength for simplicity. The single-cell model simulations predicted that the fold change in the SMAD2 N2C response at 1 h is strongly positively correlated with the amount of TGF-β receptors and weakly negatively correlated with SMAD2 and the strength of NFR (Figure 5B). However, the predicted response at 8 h was less positively correlated with the amount of TGF-β receptors and was more strongly negatively correlated with SMAD2 and NFR strength compared with the 1 h response (Figure 5C). These results suggested that heterogeneous amounts of TGF-β receptors mainly contribute to the variations of short-term SMAD signaling in single cells, whereas NFR strength and SMAD2 abundance contributes to determining long-term SMAD signaling dynamics.

**Figure 5.**
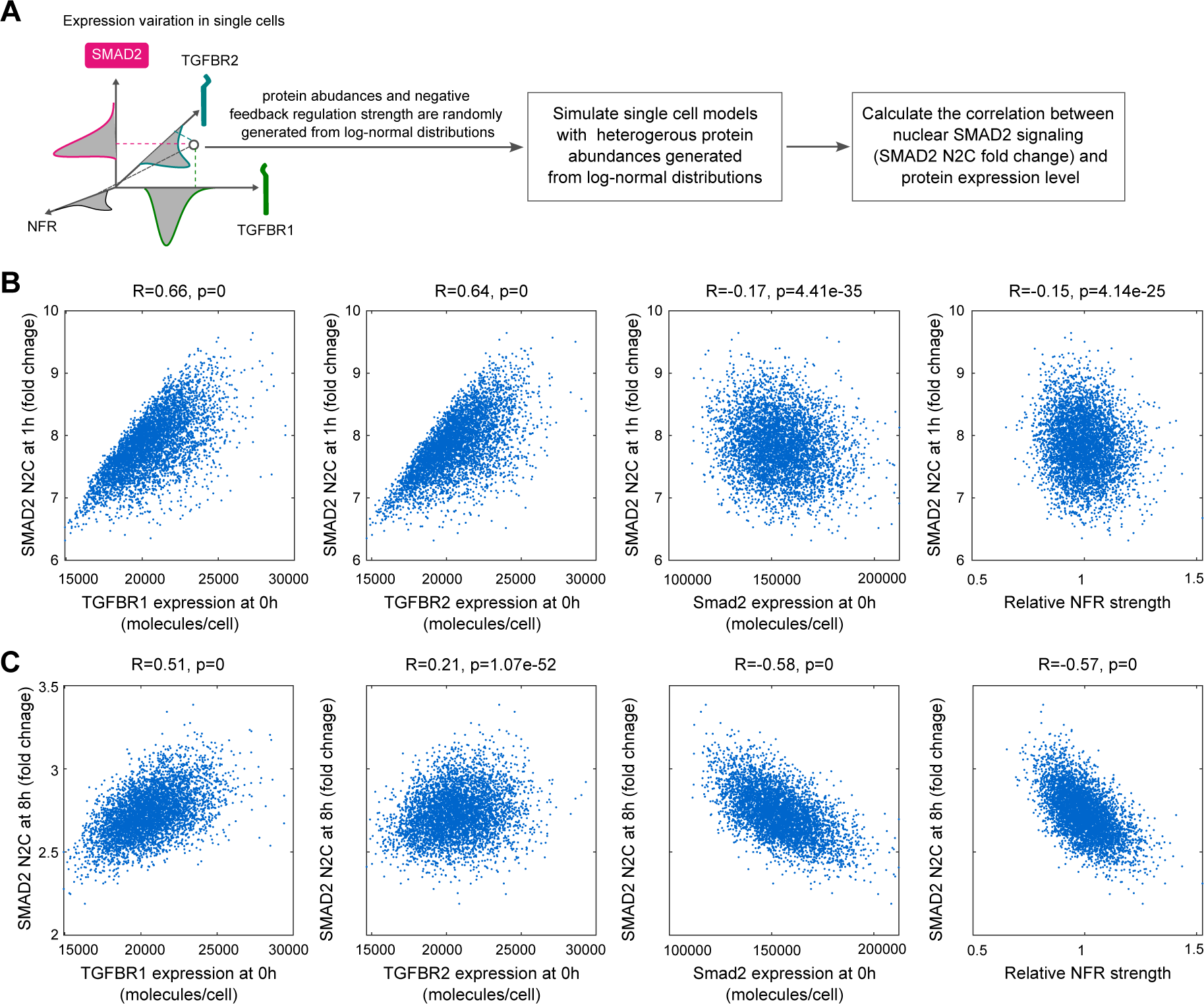
Model predictions for the correlation between the SMAD2 signaling responses and the expressions of TGF-β signaling proteins in single cells. (A) Illustration of model simulations for the SMAD2 signaling response to the variations of protein expressions in single cells. (B) Model simulations for the correlation between SMAD2 N2C fold change response at 1 h after 100 pM TGF-β stimulation and the expression of TGFBR1, TGFBR2, SMAD2 at 0 h and negative feedback regulation (NFR) strength. (C) Model simulations for the correlation between SMAD2 N2C fold change response at 8 h after 100 pM TGF-β stimulation and the expression of TGFBR1, TGFBR2, SMAD2 at 0h and negative feedback regulation (NFR) strength. The correlations (R-values and p-values) in the plots were calculated with Pearson correlation coefficients.

Using single-cell model simulations, we investigated single-cell responses in cells with imbalanced amounts of TGFBR1 and TGFBR2. The simulations were performed like those shown in Figure 5, but the mean amount of TGFBR1 or TGFBR2 was increased to 5-fold of the corresponding value in the population model. The simulations predicted that normalized nuclear SMAD2 signaling in single cells is highly correlated with the TGF-β receptor that is less abundant (Figure 6). These results resemble Liebig’s law of the minimum at a single-cell level.

**Figure 6.**
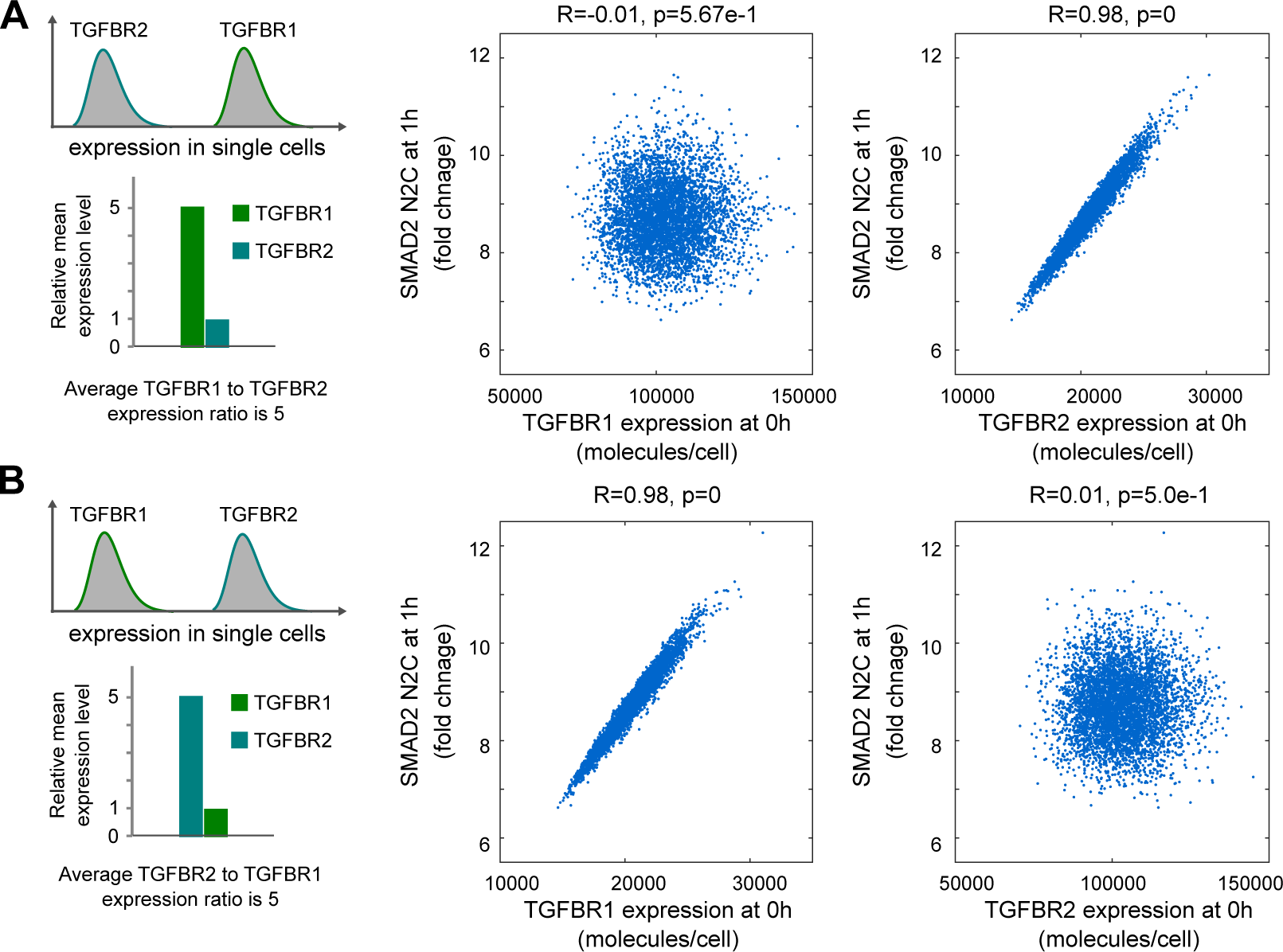
Model predictions for SMAD2 signaling responses with imbalanced expression of TGFBR1 and TGFBR2 in single cells. (A) Model predicts that SMAD2 N2C fold change is correlated with TGFBR2 when TGFBR1 is expressed at higher level than TGFBR2 in single cells (in this example, the average TGFBR1 to TGFBR2 expression ratio was set to 5). (B) Model predicts that SMAD2 N2C fold change is correlated with TGFBR1 when TGFBR1 is expressed at lower level than TGFBR2 in single cells (in this example, the average TGFBR2 to TGFBR1 expression ratio was set to 5). The correlations (R-values and p-values) in the plots were quantified with Pearson correlation coefficients.

### Experimental validation of model predictions for Liebig’s law of the minimum in single cells

Our model predictions predicted that SMAD2 signaling is correlated with both TGFBR1 and TGFBR2 when they are present in equal amounts or with constant ratios in single cells (the dashed line in Figure 3D). Therefore, we tested this prediction in the single-cell model by setting the same protein abundance values for TGFBR1 and TGFBR2 in each single cell simulation. As expected, the model predicted that SMAD2 signaling response is highly correlated with the number of TGF-β receptors in single cells (Figure 7A).

**Figure 7.**
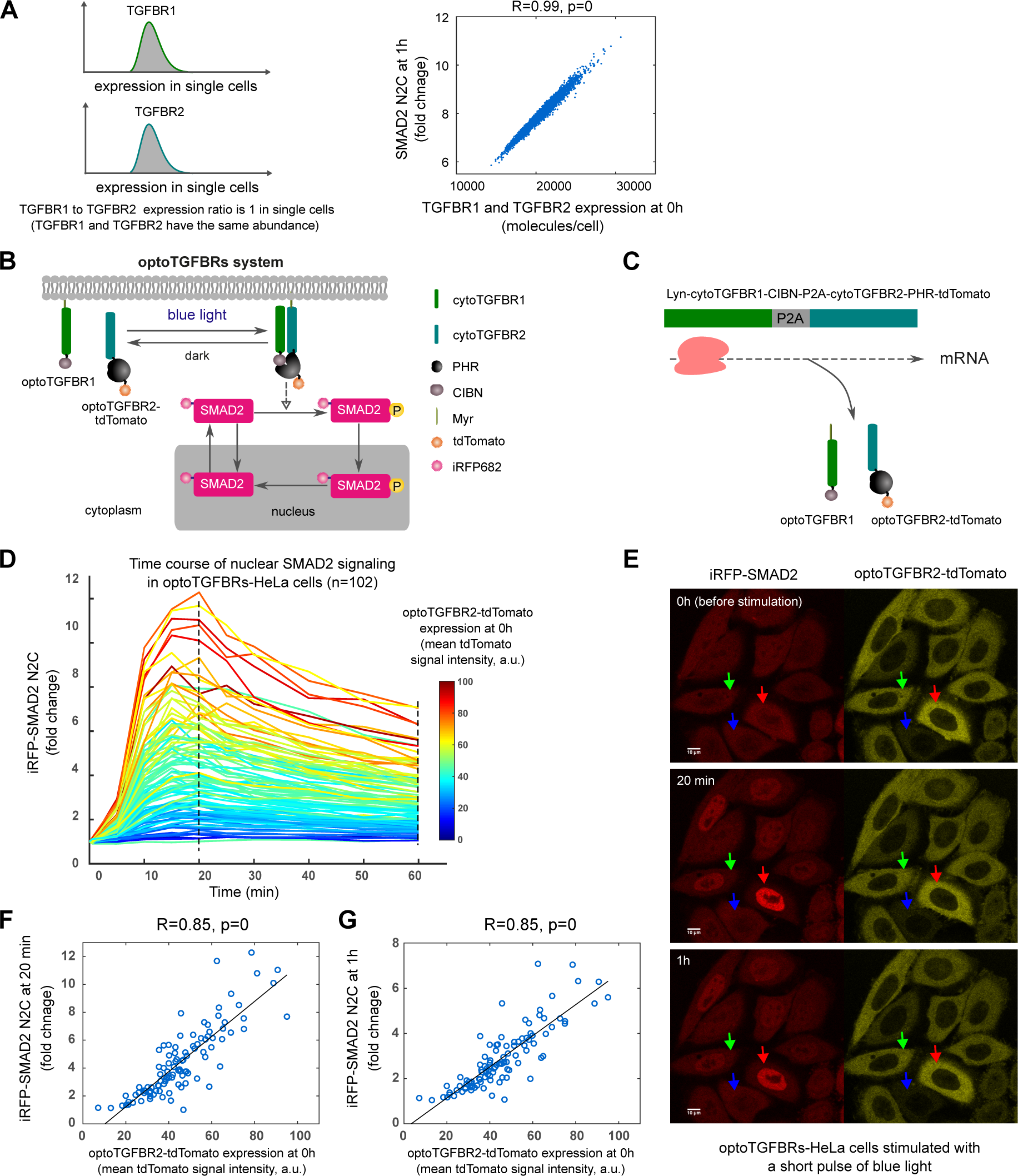
SMAD2 signaling responses are highly correlated with the amount of TGF-β receptors in single cells when TGFBR1 and TGFBR2 are expressed at a similar level. (A) Model prediction for the correlation between SMAD2 N2C fold change response at 1 h and the number of TGF-β receptors when TGFBR1 and TGFBR2 are expressed at a similar level in single cells. The protein abundance values for TGFBR1 and TGFBR2 were set to be the same in single cell simulations (TGFBR1 to TGFBR2 expression ratio is 1). (B) Schematic representation of the optoTGFBRs optogenetic system. The development of the optoTGFBRs system was described in a previous publication (Li *et al*., 2018). (C) Illustration for the co-expression of optoTGFBR1 and optoTGFBR2-tdTomato with 2A peptide element (P2A). (D) Time course profiles of iRFP-SMAD2 N2C fold change responses in individual optoTGFBRs-HeLa cells (n=102) upon one pulse of blue light stimulation. (E) Representative live-cell images of SMAD2 responses to one pulse of blue light stimulation in single cells. The red, green and blue arrows indicate cells that express optoTGFBR2-tdTomato receptor at a high, intermediate and low level, respectively. (F) The plot of iRFP-SMAD2 N2C fold change responses at 20 min versus the expression of optoTGFBR2-tdTomato before blue light stimulation (0 h) in optoTGFBRs-HeLa cells. (G) The plot of iRFP-SMAD2 N2C fold change responses at 1 h versus the expression of optoTGFBR2-tdTomato before blue light stimulation (0 h) in optoTGFBRs-HeLa cells. The correlations (R-values and p-values) in the plots were calculated with Pearson correlation coefficients.

To test these model predictions in single cells, we used the optogenetic TGF-β receptor system (optoTGFBRs) (Li *et al*, 2018). In this system, SMAD2 signaling is activated with blue light through the reversible dimerization of the N-terminal end of CIB1 (CIBN) and the PHR domain of Cryptochrome2, which are fused to chimeric TGFBR1 and TGFBR2, respectively, and fluorescent SMAD2 N2C is detected with the reporter iRFP-SMAD2 (Figure 7B). We used HeLa cells stably expressing optoTGFBR1, optoTGFBR2-tdTomato, and iRFP-SMAD2. These cells are called optoTGFBRs-HeLa. To ensure similar amounts of each receptor, a viral 2A peptide sequence (P2A) was included between the coding sequences of optoTGFBR1 and optoTGFBR2-tdTomato to induce self-cleavage during translation enabling co-expression of these two proteins (Ahier & Jarriault, 2014; Li *et al*., 2018) (Figure 7C).

With this system, we measured SMAD2 N2C fold changes in individual optoTGFBRs-HeLa cells at different time points after blue light stimulation. As predicted by the model, the amplitude of iRFP-SMAD2 signaling increased with the signal from optoTGFBR2-tdTomato, indicating the abundance of optogenetic TGFBR2 receptor, in individual cells (Figure 7D-E). Furthermore, the SMAD2 N2C fold change responses at 20 min and 1 h were highly correlated with the amount of optoTGFBR2-tdTomato (Figure 7F-G). We observed the same correlation when this system was stably expressed in HaCaT cells (optoTGFBRs-HaCaT) (Figure S12). Together, these results showed that variable but matched amounts of TGF-β receptors in individual cells produced heterogeneous TGF-β signaling responses at the single-cell level.

To further test our model predictions, we induced imbalanced expression of optoTGFBR1 and optoTGFBR2 through overexpressing optoTGFBR1 or optoTGFBR2 in optoTGFBRs-HeLa cells, which initially expressed similar levels of both receptors. Western blot analysis confirmed the desired imbalance (Figure S13). Consistent with the model predictions (Figure 6), the strong correlation between SMAD2 N2C fold change response at 1h and optoTGFBR2-tdTomato expression levels persisted in single cells when optoTGFBR1 was overexpressed (Figure 8A). Conversely, the high correlation between nuclear SMAD2 signaling and optoTGFBR2 expression levels vanished at single cell level when optoTGFBR2 was overexpressed (Figure 8B). These experimental results validate our model predictions, confirming that the SMAD2 signaling is determined by the low abundance TGF-β receptor in single cells.

**Figure 8.**
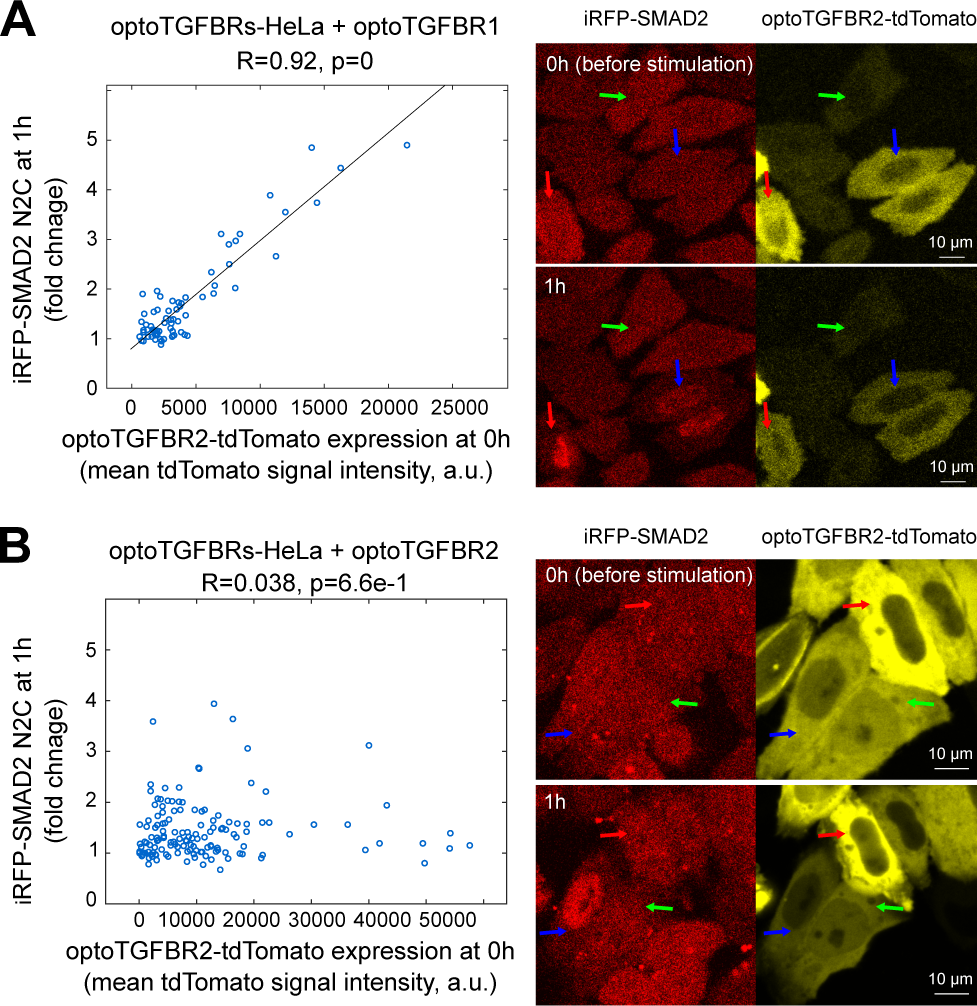
Relationship between SMAD2 signaling responses and optoTGFBR2 expression level in optoTGFBRs-HeLa cells with imbalanced expression of optoTGFBR1 and optoTGFBR2. The plot shows the fold change responses of iRFP-SMAD2 N2C at 1 h versus the expression of optoTGFBR2-tdTomato before blue light stimulation (0 h) in optoTGFBRs-HeLa cells with overexpression of optoTGFBR1 **(A)** and optoTGFBR2-tdTomato **(B)**, respectively. Representative live-cell images of SMAD2 responses to one pulse of blue light stimulation are provided on the right for each condition. The correlations (R-values and p-values) in the plots were calculated with Pearson correlation coefficients.

## Discussion

Signaling proteins are present in different quantities across cell lines and within individual cells. The variability of signaling protein abundance affects the transmission of molecular signal in a nuanced and sophisticated manner. In this study, we investigated the relationship between TGF-β signaling responses, in particular SMAD2 activity, and the variation of protein abundance. The TGF-β pathway uses two types of receptors (TGFBR1 and TGFBR2) to propagate the signal. Our analyses showed that heterogeneity in SMAD2 signaling follows specific patterns according to the variations in TGF-β receptor abundance. We found that different cell lines show distinct sensitivities to perturbation of receptor abundance depending on the relative amounts of TGFBR1 and TGFBR2. When the amounts of TGFBR1 and TGFBR2 are imbalanced, as occurs in some cell lines (such as RH-30 and HepG2 cells studied here), nuclear SMAD2 signaling is sensitive to variation of low abundance receptor but is robust to variation of the high abundance receptor (Figure 4). This feature is consistent with Liebig’s law of the minimum (Liebig, 1840). Similar to the rate-limiting concept in enzymology, our work indicated that the low abundance TGF-β receptor may serve as a rate-limiting factor that dictates the TGF-β signaling response in different cellular contexts. Similarly, we speculate that the BMP signaling pathway, utilizing two receptors for processing BMP ligands, may exhibit minority control principles at the BMP receptor level, influencing the heterogeneity of SMAD1/5/8 signaling responses. It will be of great interest to see whether the principle of minimum control governs the heterogeneity of signaling responses not only in TGF-β, but also in BMP and other signaling pathways.

The responses of TGF-β signaling can display diverse patterns in different time scales at the population and single-cell levels (Strasen *et al*., 2018; Zi *et al*., 2011). Multiple studies report that SMAD signaling in the short-term (here, considered within 1 hour) senses the amount of TGF-β through the activation of receptors (Frick *et al*., 2017; Nunns & Goentoro, 2018; Schmierer *et al*., 2008; Zi *et al*., 2011). In this work, we found that short-term SMAD signaling is highly correlated with the number of TGF-β receptors in single cells, when each receptor is present at an equivalent amount using an engineered optogenetic system (Figure 7). In the long-term (8 hours or longer), downstream TGF-β signaling patterns diverge as negative feedback regulators alter the availability of TGF-β ligand and receptors (Strasen *et al*., 2018; Zi *et al*., 2011). Our computational analysis showed that long-term SMAD signaling negatively correlated with the strength of NFR (Figure 5C). This is consistent with a study showing that knocking down SMAD7, a negative feedback regulator of TGF-β signaling, reduces the heterogeneity of long-term SMAD2 dynamics in single cells (Strasen *et al*., 2018). However, a direct quantification of the relationships between TGF-β signaling responses and feedback proteins is missing at the single-cell level. Moreover, our validation of single cell model predictions using a synthetic optogenetic TGF-β receptor system offers valuable insights, but it remains important to examine if similar results are observed in in wild-type cells stimulated with TGF-β ligand given potential differences in oligomerization and activation dynamics between synthetic and endogenous TGF-β receptors. Single-cell protein profiling approaches could provide key information about signaling relationships that produce heterogeneous responses (Gut *et al*, 2018; Kramer *et al*., 2022; Lun *et al*., 2017). Advances in this area should help to clarify the dependence of signaling responses on protein abundance in spaces with higher dimensionality.

TGF-β signaling is highly complex and regulated at multiple levels. Our current model represents a simplified version of this complexity and focuses on the core elements of the pathway, which has its limitations. The expansive TGF-β network encompasses multiple feedback loops and crosstalk with other signaling pathways (Derynck & Budi, 2019; Luo, 2017; Zhang, 2017). Incorporating mechanisms beyond our model’s current scope, such as heteromeric receptor complexes involving TGF-β and BMP receptors, SMAD linker regulation, the formation of mixed SMAD complexes and non-SMAD signaling, awaits future expansions of our model contingent on the availability of quantitative data for these processes. In addition, our model analyses used relative SMAD2 activity and nuclear SMAD2 fold change signal as readouts, exploring the effects of TGF-β receptor abundance variations on TGF-β signaling. While relative SMAD2 signaling is pertinent for comparing TGF-β signaling response patterns within the same cell type, the absolute abundance of SMAD protein could be important when comparing TGF-β signaling across different cell types. Considering that different R-SMADs and SMAD4 can form diverse homo- and hetero-oligomeric complexes, variations and mutations in SMAD abundance could modulate the balance of different SMAD complexes, thereby regulating heterogeneous gene expression in diverse cell types (Fleming *et al*, 2013; Korkut *et al*, 2018; Lucarelli *et al*., 2018). Importantly, the expression ratio of TGFBR1 and TGFBR2 within the same cell type or in single cells might dynamically change over time after exposure to stimuli or different microenvironments. Consequently, long-term TGF-β signaling responses depend not only on the initial abundance of the low abundance receptor but also on dynamic changes in the low abundance receptor, other feedback regulators (e.g., SMAD7) and non-SMAD signaling.

Our finding that the TGF-β pathway modulates variations in TGF-β receptor abundance according to Liebig’s law of the minimum raises intriguing questions about the fine-tuning of TGF-β signaling in both normal and disease contexts. Canonical TGF-β signaling conventionally involves TGFBR1 and TGFBR2 signaling through the phosphorylation of Smad2 and Smad3. However, recent studies have highlighted that TGFBR1 can interact with the type I BMP receptor ACVR1 to activate Smad1 and Smad5 in endothelial cells, and possibly during epithelial-mesenchymal transition (EMT) (Ramachandran *et al*, 2018). Similarly, ACVR2A can form stable heteromeric complexes with activin type I receptor (ALK4) and BMP type I receptors (ALK2/3/6), providing mechanism for fine-tuning two branches of Smad signaling responses to activin/BMP stimuli (Szilagyi *et al*, 2022). In instances where TGFBR1 is in low abundance, a plausible scenario arises where the high-affinity TGFBR1/ACVR1 complex competes with the TGFBR1/TGFBR2 complex for TGFBR1. This competition may result in the attenuation of canonical TGF-β signaling through Smad2/3, while TGFBR1 is engaged in TGFBR1/ACVR1 signaling through Smad1/5. Such a scenario suggests that in cancer and EMT, TGF-β signaling could undergo modulation in trans by TGFBR1/ACVR1 signaling. Expanding our current model to incorporate interactions among diverse receptors and SMADs would facilitate the exploration of design principles arising from various receptor oligomerization modes with distinct binding affinities. Nevertheless, reliable predictions rely on the comprehensive characterization of signaling protein abundances, binding affinities between ligands and receptors, and the kinetics of interactions between SMADs and receptors. Future modeling and experimental studies are necessary to elucidate how variations in TGF-β signaling in diseases are impacted when the abundances of TGFBR1 and TGFBR2 are altered.

The pivotal role of TGF-β signaling in cancer progression is evident through the frequent differential expression of genes encoding TGF-β receptors in various human cancers (Dzieran *et al*, 2013; Jakowlew *et al*, 1997; Korkut *et al*., 2018; Venkatasubbarao *et al*, 2000). Our findings underscore the significance of maintaining optimal expression levels and ratios of type I and type II receptors for effective TGF-β signaling. Partial changes in the level of the low abundance TGF-β receptor may lead to imbalanced cellular responses, thereby contributing to pathological outcomes. For instance, experimental evidence demonstrates that haploid loss of TGFBR1 and TGFBR2 impairs SMAD signaling and promote tumor progression in mouse models (Ijichi *et al*, 2006; Zeng *et al*, 2009). Many pharmacological compounds have been designed to target TGFBR1 and TGFBR2 (Colak & Ten Dijke, 2017). Our finding also indicates the importance of considering the relative abundance of TGF-β receptors for drug targeting in the TGF-β pathway. Selective inhibition of TGFBR2 could be more effective than targeting TGFBR1 in tumors with a high TGFBR1-to-TGFBR2 abundance or gene expression ratio and vice versa. Alternatively, strategies focused on upregulating the expression of low abundance TGF-β receptors could be potential therapeutic interventions to restore the beneficial function (tumor suppression) of TGF-β signaling in diseases. Given the dual roles of TGF-β in tumor progression, consideration must be given to its detrimental function in cancer metastasis. Thus, it becomes imperative to develop a comprehensive predictive model that integrates both SMAD signaling and non-SMAD signaling and elucidate how cells convert TGF-β signals into physiological functions. We envision that the principle of minimum control could help us to understand why specific signaling proteins function differently in diverse cellular contexts. Understanding the intricate relationships between signaling outputs and the abundance of signaling components holds great potential for manipulating pathway activities in therapeutic applications.

## Materials and Methods

### Cell culture and the constructs of stable cell lines

HaCaT, HepG2, optoTGFBRs-HeLa, and optoTGFBRs-HaCaT cells were cultured in DMEM medium supplemented with 10% fetal bovine serum, 100 units/mL penicillin and 100 μg/mL streptomycin, and 2 mM L-glutamine at 37 °C and 5% CO2. RH-30 cells were cultured in RPMI 1640 medium with the same supplements as those in the complete medium for culturing other cells. The optoTGFBRs-HaCaT cell line was established following the same procedure as for the development of the optoTGFBRs-HeLa cell line (Li *et al*., 2018).

### Cell lysate preparation and immunoblotting

Cell lysates were prepared with RIPA buffer (Cell Signaling, #9806), supplemented with 1 mM PMSF, 1 mM NaF, protease inhibitor cocktail (Roche, #04 693 132 001), and phosphatase inhibitor cocktail (Roche, #04 906 845 001). SDS–PAGE gels were transferred onto nitrocellulose membranes.

We used the following primary antibodies against: Phospho-SMAD2 (Ser465/467) (Cell Signaling, #3108), SMAD2 (Cell Signaling, #3103), β-Actin (Cell Signaling, #3700), TGFBR1 (Santa Cruz, #sc-398), TGFBR2 (abcam, #ab184948), and integrin β1 (Cell Signaling, #9699). All Western blot membranes were incubated with primary antibodies overnight at 4 °C. Secondary antibodies used for Western blotting were anti-rabbit IgG (H+L) DyLight 800 (Cell Signaling, #5151) and anti-mouse IgG (H+L) DyLight 680 (Cell Signaling, #5470). Western blot images were acquired and quantified using Odyssey CLx Imaging System (LI-COR Biosciences, #9140).

### siRNA-mediated knockdown

Cells were seeded in 6-well plates at 50%-70% confluency and incubated overnight. On day 2, cells were transfected with different concentrations of SMARTpool siRNAs (final concentrations: 0.08-100 nM) using DharmaFECT1 reagents following the manufacturer’s protocol. Non-targeting controls corresponded to the samples transfected with non-targeting control siRNA (final concentration: 20 nM). Each SMARTpool siRNA product contains four types of individual siRNAs targeting a single gene. After two days of transfection, cells were cultured with standard cell culture medium and where indicated the cells were stimulated with 100 pM (2.5 ng/mL) TGF-β1 for 1 h.

The following siRNA reagents were purchased from Horizon Discovery Group: TGFBR1 siRNA (catalog number: L-003929-00-0010), TGFBR2 siRNA (catalog number: L-003930-00-0010), and non-targeting control siRNA (catalog number: D-001810-10-05).

### RNA extraction and RT-qPCR analysis

HaCaT, RH30 and HepG2 cells were treated as indicated in the figures before the isolation of RNA using the RNeasy Mini Kit (Qiagen, 74104) according to the manufacturer’s instructions. The mRNA was reverse-transcribed to cDNA using the High Capacity cDNA Reverse Transcription kit reagents (Thermo Fisher Scientific, 4368813). RT–qPCR was performed in triplicates using the PowerUp SYBR Green Master Mix (Thermo Fisher Scientific, A25742) on QuantStudio 6/7 qPCR instruments. The primer sets used were list as below:

human *GAPDH*, forward: CCACTCCTCCACCTTTGAC
human *GAPDH*, reverse: ACCCTGTTGCTGTAGCCA
human *SMAD7*, forward: ACCCGATGGATTTTCTCAAACC
human *SMAD7*, reverse: GCCAGATAATTCGTTCCCCCT
human *PAI1*, forward: GAGACAGGCAGCTCGGATTC
human *PAI1* reverse: GGCCTCCCAAAGTGCATTAC
human *JUNB* forward: CAAGGTGAAGACGCTCAAGG
human *JUNB* reverse: TCATGACCTTCTGTTTGAGCTG

### Overexpression of optoTGFBR1 and optoTGFBR2 in optoTGFBRs-HeLa cells

OptoTGFBRs-HeLa cells were transiently transfected with Myr-cytTβRI-CIBN (optoTGFBR1) or cytTβRII-PHRCRY2-tdTomato plasmids (optoTGFBR2) at a concentration of 100 ng per well in 96-well plates using Lipofectamine 3000, following the manufacturer’s protocol. Transfected cells were plated on a 96-well plate and incubated for 1 day before live-cell imaging experiments. Detailed information about Myr-cytTβRI-CIBN and cytTβRII-PHRCRY2-tdTomato plasmids was previously described (Li *et al*., 2018).

### Live-cell imaging and photoactivation

Live-cell imaging experiments were performed using inverted confocal laser scanning microscopes from the Zeiss LSM series, which were equipped with incubators to maintain the samples at 37 °C and with 5% CO_2_ during the measurements.

Photoactivation experiments in optoTGFBRs-HeLa and optoTGFBRs-HaCaT cells with similar expression levels of optoTGFBR1 and optoTGFBR2 (Figure 7, Figure S12) were conducted on the LSM 710 NLO using a Plan-Apochromat 63×/1.40 oil objective. Images were acquired with a resolution of 512 x 512 pixels (pixel size 264 nm, scan area 135 x 135 µm) and a pixel dwell time of 12.6 µs. Fluorescence images (8-bit) of opto-TGFBR2-tdTomato and iRFP682-Smad2 were obtained prior to photoactivation by sequential excitation at 543 nm and 633 nm, respectively, with detection ranges from 548 to 647 nm and from 638 to 759 nm (pinholes: 152 µm and 199 µm). Laser power and gain were kept constant during experiments. The light-sensitive optogenetic system was then photoactivated by irradiation with 488-nm-laser light using the ZEN 2 software bleaching module. The activation laser power was measured with the aid of an optical power meter (Thorlabs) and was set to 12.4 µW (Li & Zi, 2022). The entire field of view was photoactivated with a pixel dwell time of 3.14 µs (scan time 0.97 s), followed by time lapse imaging of the iRFP682-Smad2 fluorescence for 60 min. (time points for optoTGFBRs-HeLa: 0.5, 5, 10, 15, 20, 25, 30, 40, 50, 60 min post-activation; time points for optoTGFBRs-HaCaT: 0.5, 2, 4, 6, 8, 10, 15, 20, 25, 30, 35, 40, 45, 50, 55, 60 min post-activation).

The optoTGFBRs-HeLa cells with imbalanced expression of optoTGFBR1 and optoTGFBR2 (Figure 8) were examined using the LSM 880 Airyscan equipped with a C-Apochromat 40×/1.2 W Korr objective. opto-TGFBR2-tdTomato and iRFP682-Smad2 were excited at 561 nm and 633 nm, respectively, with detection ranges from 566 to 638 nm and from 638 to 755 nm (pinhole: 131 µm, 3 airy units; depth of focus 3 µm). 16-bit images were acquired with a resolution of 1352 x 1352 pixels (pixel size 262 nm, scan area 354 x 354 µm) and a pixel dwell time of 6.21 µs. For photoactivation, the power of the 488-nm-laser was set to 26.3 µW, with a pixel dwell time of 12.4 µs. Images of opto-TGFBR2-tdTomato and iRFP682-Smad2 were recorded.

### Image analysis and quantification

Live-cell imaging data were manually quantified using ImageJ. We first quantified the SMAD2 N2C response by calculating the ratio of nuclear to cytoplasmic iRFP-SMAD2 signal. The SMAD2 N2C fold change response in each cell was quantified by normalizing SMAD2 N2C to the corresponding SMAD2 N2C signal at 0 h in the same cell. The optoTGFBR2-tdTomato amount was quantified as the tdTomato fluorescence intensity in the cytoplasmic area after subtracting the background signal measured in the areas without cells. Three cytoplasmic and three nuclear areas were quantified, and the average values were used for calculations. Cells that exited the field of view, underwent division, or experienced cell death were excluded from the analysis.

### Fitting of P-SMAD2 dose response curve to TGFBR1 or TGFBR2 knockdown

To quantify how P-SMAD2 responded to siRNA knockdown perturbations of TGFBR1 and TGFBR2, we fitted a Hill function to the dose response curves (Figure 4). The amount of P-SMAD2 and the amount of TGFBR1 (or TGFBR2) in each perturbation were first normalized to those in the non-targeting control (ntc) samples. Then the dose response curve was fitted with a Hill function using least squares fitting.

### Mathematical modeling

Model simulations were performed using MATLAB. Parameter estimation and identifiability analyses were performed with the parallel parameter estimation tool SBML-PET-MPI (Zi, 2011). The details about model development, assumptions, parameter estimation, the system of ordinary differential equations and kinetic parameter values are described in the Supplementary Information.

### Data availability

All data is included in the manuscript and/or supporting information.

## Acknowledgements

This work was supported by BMBF e:Bio SyBioT project (031A309) to ZZ. The authors thank Dr. Ping Wei for critical reading of this manuscript and Dr. Nancy R. Gough (BioSerendipity, LLC) for editorial support.

## Author contribution

YL, DD and ZZ: experiments and data analysis; ZZ: mathematical modeling; CTH contributed to the setup of live-cell imaging experiments; JK and WDH contributed to constructing the optoTGFBRs plasmid; QX and XL provided insights on the data analysis. ZZ wrote the manuscript with contributions from all the authors. ZZ conceived the study and supervised the research.

