## Supplementary Material for "Liebig’s law of the minimum in the TGF-β/SMAD pathway"

### 1. Supplementary Methods

#### *1.1 The minimal mathematical model of the TGF-β pathway*

To investigate the signaling activity profiles of the TGF-β pathway in response to variations in TGF-β receptors, we developed a minimal model that captures the essential components of the pathway through two reactions. The first reaction represents the binding of the TGF-β ligand ($L$) to type I ($R_1$) and type II ($R_2$) TGF-β receptors, forming the ligand-receptor complex ($LRC$). The second reaction describes the phosphorylation and dephosphorylation of the SMAD protein ($S_u$: unphosphorylated Smad, $S_p$: phosphorylated Smad).

Following the approach used in published BMP pathway models (Antebi *et al*, 2017; Su *et al*, 2022), we model the binding and unbinding of TGF-β ligand and its receptors as one-step reversible reaction with forward rate ($k_f$) and backward rate ($k_b$). We assumed that the phosphorylation of the SMAD protein is proportional to the activity of the ligand-receptor complex. In this minimal model, our focus is on the steady-state signaling activity of the TGF-β pathway. Additionally, we assumed the total amounts of receptors and SMAD proteins to remain constant (conservation of mass), following similar assumptions made in mathematical models of TGF-β and BMP pathway (Antebi *et al.*, 2017; Su *et al.*, 2022; Vilar *et al*, 2006). The two reactions can be summarized as:

$$L + R_1 + R_2 \underset{k_b}{\overset{k_f}{\rightleftarrows}} LRC \tag{1}$$

$$S_u \underset{k_{depho}}{\overset{k_{pho}[LRC]}{\rightleftarrows}} S_p \tag{2}$$

where $S_u$ denotes unphosphorylated SMAD, $S_p$ denotes phosphorylated SMAD.

Based on mass-action kinetics, we can formulate the following set of ordinary differential equations to describe the dynamics of these reactions:

$$\frac{d[R_1]}{dt} = k_b[LRC] - k_f[L][R_1][R_2] \tag{3}$$

$$\frac{d[R_2]}{dt} = k_b[LRC] - k_f[L][R_1][R_2] \tag{4}$$

$$\frac{d[LRC]}{dt} = k_f[L][R_1][R_2] - k_b[LRC] \tag{5}$$

$$\frac{d[pS]}{dt} = k_{pho}[S_u][LRC] - k_{depho}[S_p] \tag{6}$$

With the mass of conservation, we obtain the following constraints:

$$R_1^t = [R_1] + [LRC] \tag{7}$$

$$R_2^t = [R_2] + [LRC] \tag{8}$$

$$S^t = [S_u] + [S_p] \tag{9}$$

where the variables with superscript '*t*' denote the total of the respective species with mass conservation.

Furthermore, in typical experimental setups, the TGF-β ligand is prepared in a much larger volume of media compared to the volume of cells, resulting in a scenario where there are significantly more ligand molecules than receptors. In such cases, the concentration of the TGF-β ligand remains constant.

$$[L] = L^t \tag{10}$$

By plugging equation 9 to equation 6 and scaling it to $S^t$, equation 6 can be rewritten as:

$$\frac{dS_a}{dt} = k_{pho}(1 - S_a)[LRC] - k_{depho} \cdot S_a \tag{11}$$

where $S_a$ represents the TGF-β signaling activity, which is the percentage of total SMAD proteins that are actively phosphorylated.

We next focused on the TGF-β signaling activity ($S_a$) at steady state after ligand stimulation, which can be solved based on the following algebraic equations:

$$k_b[LRC^{ss}] - k_f[L^{ss}][R_1^{ss}][R_2^{ss}] = 0 \tag{12}$$

$$k_{pho}(1 - S_a^{ss})[LRC^{ss}] - k_{depho} \cdot S_a^{ss} = 0 \tag{13}$$

Plugging equations 7-9 to equations 12 and 13, we can derive the steady state of the TGF-β signaling activity ($S_a^{ss}$) as the following:

$$LRC^{ss} = \frac{\left(1+\frac{k_f}{k_b}L^tR_1^tR_2^t\right) - \sqrt{\left(1+\frac{k_f}{k_b}L^tR_1^tR_2^t\right)^2 - 4\left(\frac{k_f}{k_b}L^t\right)^2 R_1^tR_2^t}}{2\frac{kf}{kb}L^t} \tag{14}$$

$$S_a^{ss} = \frac{k_{pho}[LRC^{ss}]}{k_{pho}[LRC^{ss}] + k_{depho}} \tag{15}$$

Defining the following parameters

$$K_1 = \frac{k_f}{k_b} \tag{16}$$

$$K_2 = \frac{k_{pho}}{k_{depho}} \tag{17}$$

$$\alpha = 1 + K_1 L^t R_1^t R_2^t \tag{18}$$

$$\beta = 4\left(K_1 L^t\right)^2 R_1^t R_2^t \tag{19}$$

Equation 14 and 15 can be rewritten as

$$[LRC^{ss}] = \frac{\alpha - \sqrt{\alpha^2 - \beta}}{2K_1 L^t} \tag{20}$$

$$S_a^{ss} = \frac{K_2[LRC^{ss}]}{K_2[LRC^{ss}]+1} = \frac{K_2\left(\alpha - \sqrt{\alpha^2-\beta}\right)}{K_2\left(\alpha-\sqrt{\alpha^2-\beta}\right)+2K_1L^t} \tag{21}$$

With equation 18, 19 and 21, we can plot the response profiles of the TGF-β signaling activity at steady state ($S_a^{ss}$) with the combinations of different R1 and R2 values. The initial abundance ($N_X$, molecules per cell) and the initial concentration ($X^0$) of specie $X$ can be exchanged with the following equation:

$$N_X = N_A V_{cell} X^0 \tag{22}$$

where $N_A$ is Avogadro number (6.02×1023). $V_{cell}$ is set to be 3.3 pL (3.3×10-12 L) according to previous reports (Schmierer et al, 2008). It's worth noting that the choice of $V_{cell}$ value does not change the response profiles shown in Figure 2 and Figure S1 as cell volume is just a scaling factor for the conversion between specie concentration and abundance.

##### *1.2 The extended model of the TGF-β pathway*

We next developed an extended mathematical model based on our previously published TGF-β model (Zi *et al*, 2011). Compared to the minimal model, the extended model includes additional TGF-β signaling processes: (1) receptor production, degradation, endocytosis and activation; (2) SMAD2 nucleocytoplasmic shuttling, phosphorylation and dephosphorylation; (3) negative feedback on the degradation of ligand receptor complex; (4) reversible binding of TGF-β to the cell surface. In the extended model, we used a linear chain of 2 variables (z1, z2) with linear differential equations to model the time delay between nuclear SMAD2 activity (PSmad2n) and the negative feedback on the degradation of ligand-receptor complex.

The extended model consists of 13 species and 22 reactions, characterized by 20 kinetic parameters. Details regarding the initial conditions, parameter values, and equations can be found in Table S2-S5. To determine the initial condition of the extended model, we measured the abundance of SMAD2 and TGF-β receptors in the HaCaT cell line (as described in section 1.2). Additionally, we derived 11 kinetic parameters from experimental datasets, including our own and those from other studies. The remaining 9 unknown parameter values were estimated by fitting the model to 102 experimental data points obtained from HaCaT cells exposed to different TGF-β stimulation conditions. Notably, parameter identifiability analysis confirms the identifiability of these parameters (see section 1.3.5 for more details on the estimation of unknown parameters).

##### *1.3 Parameter estimation for the extend model*

###### *1.3.1 The abundance of SMAD2 and TGF-β receptors*

To determine the abundance of SMAD2 and TGF-β type II receptor (TGFBR2, referred to as T2R in the extended model), we performed quantitative immunoblotting on HaCaT cells. This technique allowed us to estimate the absolute protein levels based on a standard curve signal generated from recombinant protein standards. The detailed procedure is described in a published protocol (Deng & Zi, 2022). The measured abundances of SMAD2 and TGFBR2 proteins in HaCaT are 1.54e5 and 2.06e4 molecules/cell, respectively (Figure S2).

According to the Human Protein Atlas database, the expression of TGFBR1 RNA in HaCaT cells is similar to that of TGFBR2 (Figure 3E). In the extended model, we set the protein abundance of TGFBR1 to be the same as TGFBR2 (2.06e4 molecules/cell), which aligns with a previously reported value (Wakefield et al, 1987).

###### *1.3.2 TGF-β receptor internalization and recycling rate constants*

We measured the internalization rate of TGFBR2 using a reversible biotinylation method (Gabriel *et al*, 2009). We next fitted the relative change of surface receptor protein over time with an exponential decay function and successfully measured the internalization rate constant of TGFBR2: $k_i = 0.022\ min^{-1} = 1.32\ h^{-1}$ (Figure S3).

It is known that most of TGF-β receptors are internalized in endosomes and about 10% of TGF-β receptors locate at cell surface (Di Guglielmo *et al*, 2003). Here we set the recycling rate is 1/9 of the internalization rate, $k_r = k_i/9 = 0.147\ h^{-1}$, this will result in about 10% receptors at cell surface in our model.

*1.3.3 TGF-β receptor production and degradation rate constants*

We measured the half-lives of TGF-β receptors and SMAD2 proteins in HaCaT cells by cycloheximide chase assays (Kao *et al*, 2015). The measured half-lives for TGFBR1, TGFBR2 and SMAD2 are 6.6 h, 3.67h, >24 h, respectively. The SMAD2 protein is stable before and after TGF-β stimulation. It is reasonable to assume the total amount of SMAD2 protein is constant in the model.

By rescaling the distribution of internalized receptors in endosomes, we can derive the degradation rate constants for T1R: $k_{deg}^{T1R} = 0.117\ h^{-1}$ ($\frac{-ln0.5}{6.6h\times0.9}$), for T2R: $k_{deg}^{T2R} = 0.21\ h^{-1}$ ($\frac{-ln0.5}{3.67h\times0.9}$). Since TGF-β ligand receptor complexes have similar distribution and half-life as the T1R receptor at resting state (Di Guglielmo *et al.*, 2003), we set $k_{deg}^{LCR} = k_{deg}^{T1R} = 0.117\ h^{-1}$.

The initial conditions of TGF-β receptors are determined by the receptor production, internalization, recycling and degradation rate constants (Zi *et al.*, 2011; Zi & Klipp, 2007). At resting state, we derive the following equations for TGF-β receptors at cell surface (*T1R$_s$*, *T2R$_s$*) and in cell cytosol (*T1R$_c$*, *T2R$_c$*):

$$[T1R_s]_0 = \frac{k_{prod}^{T1R}\cdot kr}{k_{deg}^{T1R}\cdot ki} \quad (23)$$

$$[T1R_c]_0 = \frac{k_{prod}^{T1R}}{k_{deg}^{T1R}} \quad (24)$$

$$N_{T1R} = V_{cyt}([T1R_s]_0 + [T1R_c]_0)N_A \quad (25)$$

$$[T2R_s]_0 = \frac{k_{prod}^{T2R}\cdot kr}{k_{deg}^{T2R}\cdot ki} \quad (26)$$

$$[T2R_c]_0 = \frac{k_{prod}^{T2R}}{k_{deg}^{T2R}} \quad (27)$$

$$N_{T2R} = V_{cyt}([T2R_s]_0 + [T2R_c]_0)N_A \quad (28)$$

where $N_A$ is Avogadro number ($6.02\times10^{23}$), $N_{T1R}$ and $N_{T2R}$ are the abundances of TGFBR1 and TGFBR2 in molecules/cell, respectively.

The volume of cytoplasm ($V_{cyt}$) and nucleus ($V_{nuc}$) are set as 2.3 pL ($2.3\times10^{-12}$ L) and 1 pL ($1\times10^{-12}$ L), respectively according to previous reports (Schmierer *et al.*, 2008). Solving equation 1-6, we can derive the initial conditions for T1R and T2R at cell surface and in cytosol with the following equations:

$$[T1R_s]_0 = \frac{N_{T1R}}{V_{cyt}N_A} \cdot \frac{kr}{ki+kr} = 1.49 \times 10^{-9}\, M = 1.49\, nM \tag{29}$$

$$[T1R_c]_0 = \frac{N_{T1R}}{V_{cyt}N_A} \cdot \frac{ki}{ki+kr} = 13.4 \times 10^{-9}\, M = 13.4\, nM \tag{30}$$

$$[T2R_s]_0 = \frac{N_{T2R}}{V_{cyt}N_A} \cdot \frac{kr}{ki+kr} = 1.49 \times 10^{-9}\, M = 1.49\, nM \tag{31}$$

$$[T2R_c]_0 = \frac{N_{T2R}}{V_{cyt}N_A} \cdot \frac{ki}{ki+kr} = 13.4 \times 10^{-9}\, M = 13.4\, nM \tag{32}$$

Based on the equations 24, 30 and equations 27, 32, we derived the corresponding production rate constant for T1R ant T2R:

$$k_{prod}^{T1R} = k_{deg}^{T1R}[T1R_c]_0 = 0.117 \times 13.4 = 1.57\, nM/h \tag{33}$$

$$k_{prod}^{T2R} = k_{deg}^{T2R}[T2R_c]_0 = 0.21 \times 13.4 = 2.81\, nM/h \tag{34}$$

*1.3.4 Initial conditions for cytosol and nuclear SMAD2*

SMAD2 nuclear import and export rate constants has been experimentally measured (Schmierer & Hill, 2005; Schmierer *et al.*, 2008): $k_{imp}^{Smad2} = 0.156\, min^{-1} = 9.36\, h^{-1}$ , $k_{exp}^{Smad2} = 0.763\, min^{-1} = 45.8\, h^{-1}$. We set the distribution of SMAD2 in cytoplasm and nucleus in steady state before TGF-β stimulation with the following equations

$$[Smad2_c]_0 = \frac{N_{Smad2}}{V_{cyt}N_A} \cdot \frac{k_{exp}^{Smad2}}{k_{imp}^{Smad2} + k_{exp}^{Smad2}} = 92.4\, nM \tag{35}$$

$$[Smad2_n]_0 = \frac{N_{Smad2}}{V_{nuc}N_A} \cdot \frac{k_{imp}^{Smad2}}{k_{imp}^{Smad2} + k_{exp}^{Smad2}} = 43.4\, nM \tag{36}$$

where $N_A$ is Avogadro number ($6.02\times10^{23}$), $N_{SMAD2}$ is the abundances of SMAD2 in molecules/cell.

*1.3.5 Estimation of the unknown parameters*

We measured or derived most parameter values in the refined models based on experimental data. The detailed values for these parameters are shown in Table S2. In the refined model, there were still some parameters that cannot be measured or derived directly from experimental data. For estimation of these unknown parameters, we used a parallel parameter estimation tool SBML-PET-MPI (Zi, 2011), which includes a global optimization algorithm (SRES) using least squares to minimize the sum of squares of differences between model simulations and the corresponding experimental datasets. We estimated the values of 9 parameters in the model with 102 experimental data points collected from HaCaT cells with different TGF-β stimulation conditions, which includes.

(1) Time course of medium TGF-β (TGFb_ex in the model) and SMAD2 phosphorylation (P-SMAD2) levels in response to 10 pM, 100 pM and 400pM TGF-β stimulations.

(2) Time course of cytosol and nuclear SMAD2 in response to 100 pM TGF-β stimulation.

(3) Relative P-SMAD2 level at 45 min in responses to different doses of TGF-β stimulations.

After parameter optimization, the refined model can reproduce various experimental data with different doses of TGF-β stimulations (Figure S4). To check the identifiability of the estimated parameters in the model, we performed profile likelihood analysis (Raue *et al*, 2009) and calculate 95% confidence intervals for the estimated parameters in the model. The profile likelihood analysis indicate that the estimated parameters are practically identifiable with the experimental datasets used for parameter estimation.

##### *1.4. Single cell model simulations*

To simulate the signaling responses in single cells, we use the same model as the population model except that the abundances of TGFBR1 ($N_{T1R}$), TGFBR2 ($N_{T2R}$) and SMAD2 ($N_{Smad2}$) are randomly generated from log-normal distributions with a mean of the corresponding reference value in the population model and a CV of 0.1. For each single cell model, we adjust the receptor production parameter values, while keep other kinetic parameter values fixed. The production rate constants for TGFBR1 (kprd_T1R) and TGFBR2 (kprd_T2R) are set according to equation 33 and equation 34, respectively. The initial conditions for surface and cytosol receptors can be derived with equation 29-32. The initial concentrations of cytosol SMAD2 and nuclear SMAD2 were calculated based on equation 35-36 accordingly. For the variation of negative feedback regulation (NFR), we randomly generate a value for k_NFR from a log-normal distribution with a mean of the corresponding reference value in the population model and a CV of 0.1. For each case, we performed 5000 single cell model simulations and the correlation analysis results were calculated with Pearson correlation coefficients.

##### *1.5 Measurement of TGF-β concentration in media*

Medium supernatants were collected from the culture of HaCaT cells at different time points after TGF-β1 (R&D Systems, Catalog Number: 240-B-002) stimulations. Medium TGF-β1 concentrations were measured using a TGF beta 1 ELISA kit (invitrogen, Catalog Number: 88-8350) following the manufacturer's instructions.

##### *1.6 Cell surface protein biotinylation and internalization*

The internalization rate of TGFBR2 was determined using a reversible biotinylation method (Gabriel *et al.*, 2009) with a cell surface protein isolation kit (Thermo Scientific Pierce, Catalog Number: 89881). Cell surface proteins were labeled with biotinylation reagent (sulfo-NHS-SS-biotin) at a low temperature (4 °C). A portion of the cells were shifted to permissive conditions for trafficking at 37 °C, allowing the biotinylated receptor proteins to internalize. Another set of cells was maintained at the low temperature as controls for total surface protein at time=0 and for stripping control. After different time of internalization, cells were moved back to low temperature to stop internalization and the residual surface biotin was stripped off with a reducing agent, which cleaves the disulfide-coupled biotin. The internalized receptor proteins with biotin were protected and they were not stripped. After cell lysis, biotinylated proteins were pulled down by streptavidin affinity chromatography according to the manufacturer's protocol and they were measured using quantitative immunoblotting. By subtracting the internalized receptor protein from the total surface protein control at time=0, the remaining surface receptor protein at different time points was determined. The dynamics of the remaining surface TGFBR2 were fitted with an exponential decay function to estimate the internalization rate constant.

#### 2. Supplementary Tables

##### Table S1: Observation of imbalanced *TGFBR1* and *TGFBR2* expression levels in HepG2 and RH30 cell lines across different RNA-Seq databases

| Cell Line | TGFBR1 | TGFBR2 | TGFBR1-to-TGFBR2 ratio | Data Source |
|---|---|---|---|---|
| HepG2 | 2.94 | 48.46 | 0.06 | ENCODE |
| HepG2 | 5.05 | 58.49 | 0.09 | Cell Model Passports (Sanger) |
| HepG2 | 15.9 | 114 | 0.14 | Human Protein Atlas |
| RH30 | 32.96 | 9.51 | 3.47 | Cell Model Passports (Sanger) |
| RH30 | 171.4 | 40.6 | 4.22 | Human Protein Atlas |

Note: RNA expression levels are presented in normalized transcripts per million (nTPM) for data from Human Protein Atlas and in fragments per kilobase per million mapped fragments (FPKM) for data from other databases.

**Table S2: Summary of the derived parameter values based on experimental data**

| Parameter | Value and Unit | Annotation | Reference |
|---|---|---|---|
| ki | 1.32 $h^{-1}$ | receptor internalization rate constant | this work |
| kr | 0.147 $h^{-1}$ | receptor recycling rate constant | this work |
| kprod_T1R | 1.57 nM/h | TGFBR1 production rate constant | this work |
| kprod_T2R | 2.81 nM/h | TGFBR2 production rate constant | this work |
| kdeg_T1R | 0.117 $h^{-1}$ | constitutive degradation rate constant for TGFBR1 | this work |
| kdeg_T2R | 0.21 $h^{-1}$ | constitutive degradation rate constant for TGFBR2 | this work |
| kdeg_LRC | 0.117 $h^{-1}$ | constitutive degradation rate constant for ligand receptor complex (LRC) | this work |
| kdeg1_TGFb | 20.8 $h^{-1}$ | constitutive degradation rate constant for internalized TGF-β | (Kaminska *et al*, 2005; Wakefield *et al*, 1990) |
| kimp_Smad2 | 9.36 $h^{-1}$ | SMAD2 nuclear import rate constant | (Schmierer *et al.*, 2008; Zi *et al.*, 2011) |
| kexp_Smad2 | 45.8 $h^{-1}$ | SMAD2 nuclear export rate constant | (Schmierer *et al.*, 2008; Zi *et al.*, 2011) |
| kimp_PSmad2 | 53.3 $h^{-1}$ | P-SMAD2 nuclear import rate constant | (Schmierer *et al.*, 2008) |
| Vcyt | 2.3$e^{-12}$ l | Cytoplasm volume | (Schmierer *et al.*, 2008) |
| Vnuc | 1$e^{-12}$ l | Nucleus volume | (Schmierer *et al.*, 2008) |
| Vmed | 2$e^{-9}$ l | Extracellular space volume | (Zi *et al.*, 2011) |

Note: The derived parameters were rounded with 3 effective digits

#### Table S3: Statistical analysis of the estimated parameters for the extended model

| Parameter (Unit) | Best fit | 95% confidence interval | Annotation |
|---|---|---|---|
| ka_LRC ($nM^{-2}h^{-1}$) | 6.57019e+02 | (6.25195e+02; 6.89870e+02) | Activation rate constant for ligand-receptor complex |
| kdiss_LRC ($h^{-1}$) | 1.74990e+00 | (1.68428e+00; 1.82099e+00) | Dissociation rate constant for ligand-receptor complex |
| k_NFR ($nM^{-1}h^{-1}$) | 6.04124e-02 | (5.89965e-02; 6.18755e-02) | Rate constant for negative feedback regulation |
| kdeg2_TGFb ($h^{-1}$) | 1.38784e-01 | (1.25339e-01; 1.52663e-01) | Degradation of TGF-β through non-specific binding |
| kpho_Smad2 ($nM^{-1}h^{-1}$) | 1.68603e+00 | (1.64520e+00; 1.72687e+00) | SMAD2 phosphorylation rate constant |
| kdepho_Smad2 ($h^{-1}$) | 1.94430e+00 | (1.89266e+00; 1.99747e+00) | SMAD2 dephosphorylation rate constant |
| kon_ns ($h^{-1}$) | 3.18848e-01 | (3.09881e-01; 3.27816e-01) | Association rate constant for non-specific binding of TGF-β |
| koff_ns ($h^{-1}$) | 1.00995e+00 | (9.73658e-01; 1.04783e+00) | Dissociation rate constant for non-specific binding of TGF-β |
| tau (h) | 2.00973e+00 | (1.95320e+00; 2.06782e+00) | Delay time between negative feedback regulation and P-Smad2n |

Note: The estimated parameters were rounded with 6 effective digits.

#### Table S4: Initial conditions of the extended model

| Species | Initial Conditions (Unit: nM) | Annotation |
|---|---|---|
| TGFb_ex | to be specified | TGF-β concentration in the medium |
| T1Rs | 1.49 | type I receptor at plasma membrane (cell surface) |
| T1Rc | 13.4 | type I receptor in early endosome |
| T2Rs | 1.49 | type II receptor at plasma membrane (cell surface) |
| T2Rc | 13.4 | type II receptor in early endosome |
| LRCs | 0 | LRC at plasma membrane (cell surface) |
| LRCc | 0 | LRC in early endosome |
| Smad2c | 92.4 | SMAD2 in cytoplasm |
| Smad2n | 43.4 | SMAD2 in nucleus |
| PSmad2c | 0 | phosphorylated SMAD2 monomer in cytoplasm |
| PSmad2n | 0 | phosphorylated SMAD2 monomer in nucleus |
| TGFb_c | 0 | TGF-β in early endosome |
| TGFb_ns | 0 | non-specific bound TGF-β |
| z1 | 0 | linear chain variable for the delay between PSmad2n and negative feedback regulation (NFR) |
| z2 | 0 | linear chain variable for the delay between PSmad2n and negative feedback regulation (NFR) |

#### Table S5: Ordinary differential equations of the extended model

d[TGFb_ex]/dt = koff_ns*[TGFb_c] - kon_ns*[TGFb_ex] -
Vcyt*kass_LRC*[TGFb_ex]*[T1Rs]*[T2Rs]/Vmed
d[T1Rs]/dt = kr*[T1Rc] - ki*[T1Rs] - kass_LRC*[TGFb_ex]*[T1Rs]*[T2Rs]
d[T1Rc]/dt = kprd_T1R - kdeg_T1R*[T1Rc] + kdiss_LRC*[LRCc] + ki*[T1Rs] - kr*[T1Rc]
d[T2Rs]/dt = kr*[T2Rc] - ki*[T2Rs] - kass_LRC*[TGFb_ex]*[T1Rs]*[T2Rs]
d[T2Rc]/dt = kprd_T2R - kdeg_T2R*[T2Rc] + kdiss_LRC*[LRCc] + ki*[T2Rs] - kr*[T2Rc]
d[LRCs]/dt = kass_LRC*[TGFb_ex]*[T1Rs]*[T2Rs] - k_NFR*[z2]*[LRCs] - ki*[LRCs]
d[LRCc]/dt = ki*[LRCs] - kdiss_LRC*[LRCc] - kdeg_LRC*[LRCc]
d[Smad2c]/dt = Vnuc*kexp_Smad2*[Smad2n]/Vcyt - kpho_Smad2*([LRCs]+[LRCc])*[Smad2c] -
kimp_Smad2*[Smad2c]
d[Smad2n]/dt = Vcyt*kimp_Smad2*[Smad2c]/Vnuc + kdepho_Smad2*[PSmad2n] - kexp_Smad2*[Smad2n]
d[PSmad2c]/dt = kpho_Smad2*([LRCs]+[LRCc])*[Smad2c] - kimp_PSmad2*[PSmad2c]
d[PSmad2n]/dt = Vcyt*kimp_PSmad2*[PSmad2c]/Vnuc - kdepho_Smad2*[PSmad2n]
d[TGFb_c]/dt = kon_ns*[TGFb_ex] - koff_ns*[TGFb_c] - kdeg2_TGFb*[TGFb_c]
d[TGFb_ns]/dt = kdiss_LRC*[LRCc] - kdeg1_TGFb*[TGFb_ns]
d[z1]/dt = (2*([PSmad2n] - [z1]))/tau
d[z2]/dt = (2*([z1] - [z2]))/tau

#### 3. Supplementary Figures

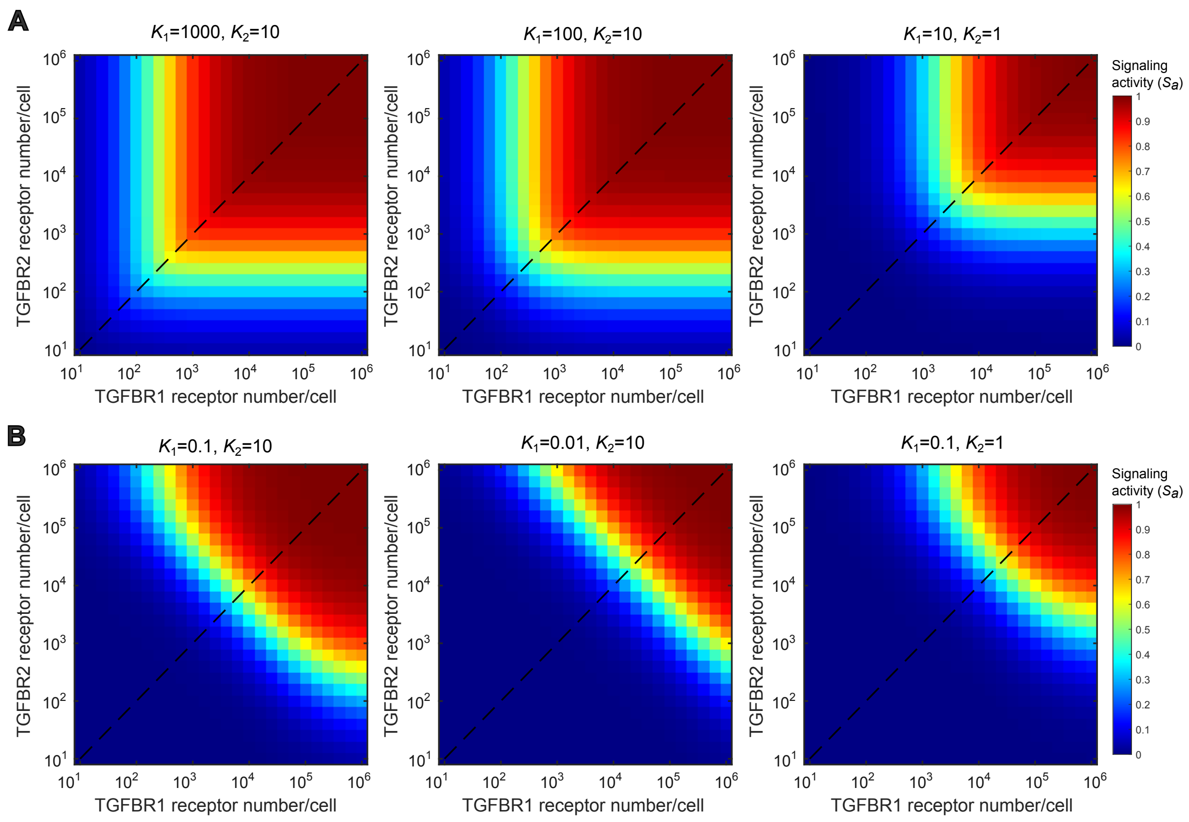

**Figure S1. The predicted response profile of TGF-β signaling activities ($S_a$) at steady state with different choice of parameter values in the minimal model.**

(A) Simulations with high binding affinity values for $K_1$ ($K_1 = 1000$, 100, 10).

(B) Simulation with low binding affinity values for $K_1$ ($K_1 = 0.1$, 0.01).

The ligand concentration $L$ is set to be 0.1 nM (100 pM) in all the simulations.

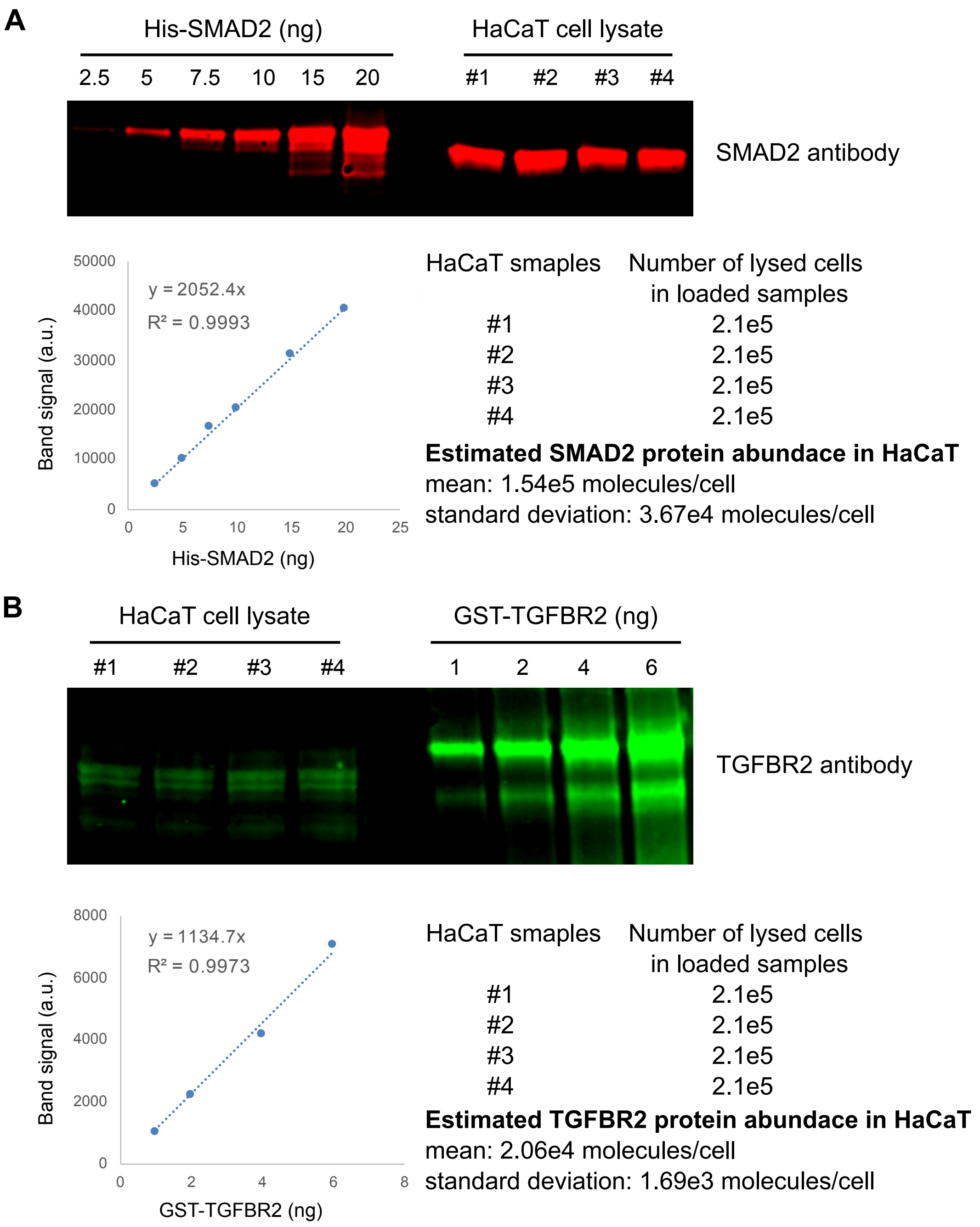

**Figure S2. Measurement of protein abundance in HaCaT cells**

Protein abundances were determined by quantitative immunoblotting with a series of recombinant protein standard.

(A) Estimation of SMAD2 abundance in HaCaT cells.

(B) Estimation of TGFBR2 abundance in HaCaT cells.

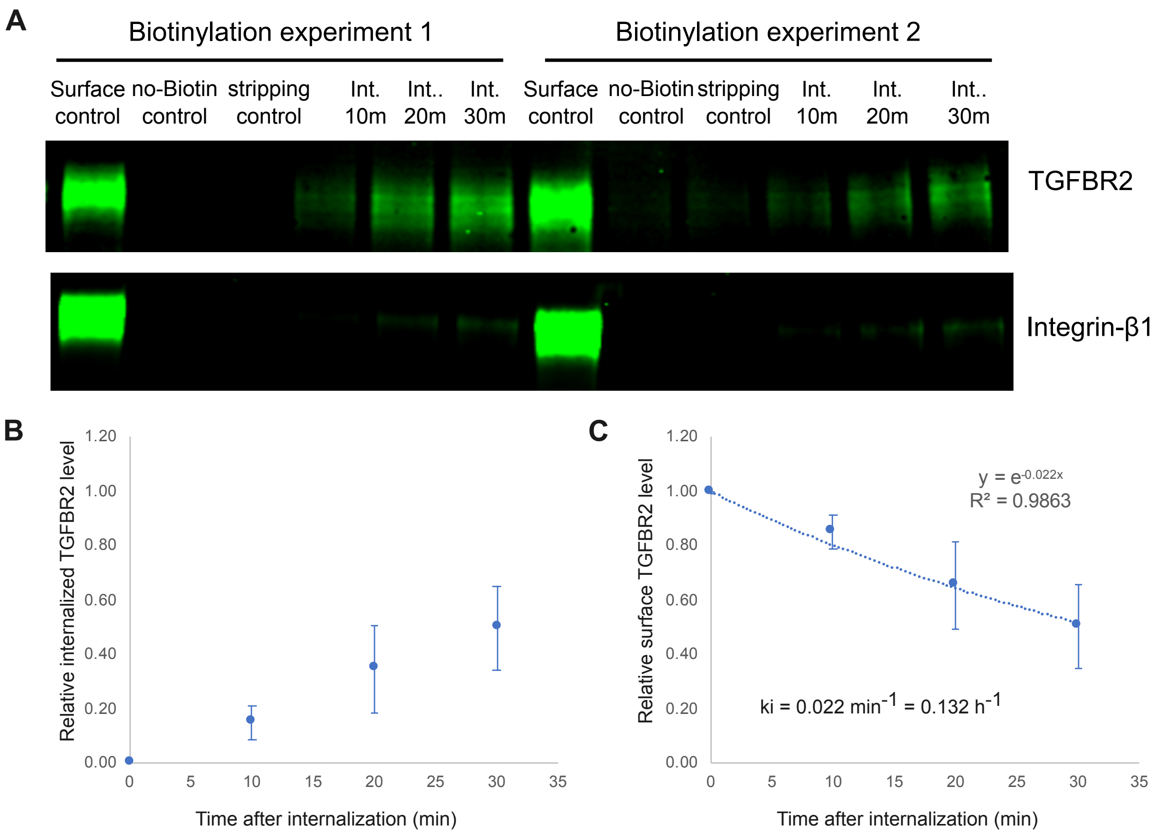

**Figure S3. Measurement of TGFBR2 internalization rate constant in HaCaT cells**

(A) Immunoblotting of the biotinylated protein samples. The biotin labeling procedure is described in the Supplementary Methods 1.6 section. Integrin-β1 is a cell surface protein marker.

(B) The kinetics of the internalized TGFBR2 were quantified from Western blots. The mean values and standard deviations from 4 experiments are shown.

(C) The kinetics of the remaining surface TGFBR2 after internalization. The mean values and standard deviations from 4 experiments are shown. The data points were fitted to an exponential decay function and the internalization rate constant (ki) was estimated.

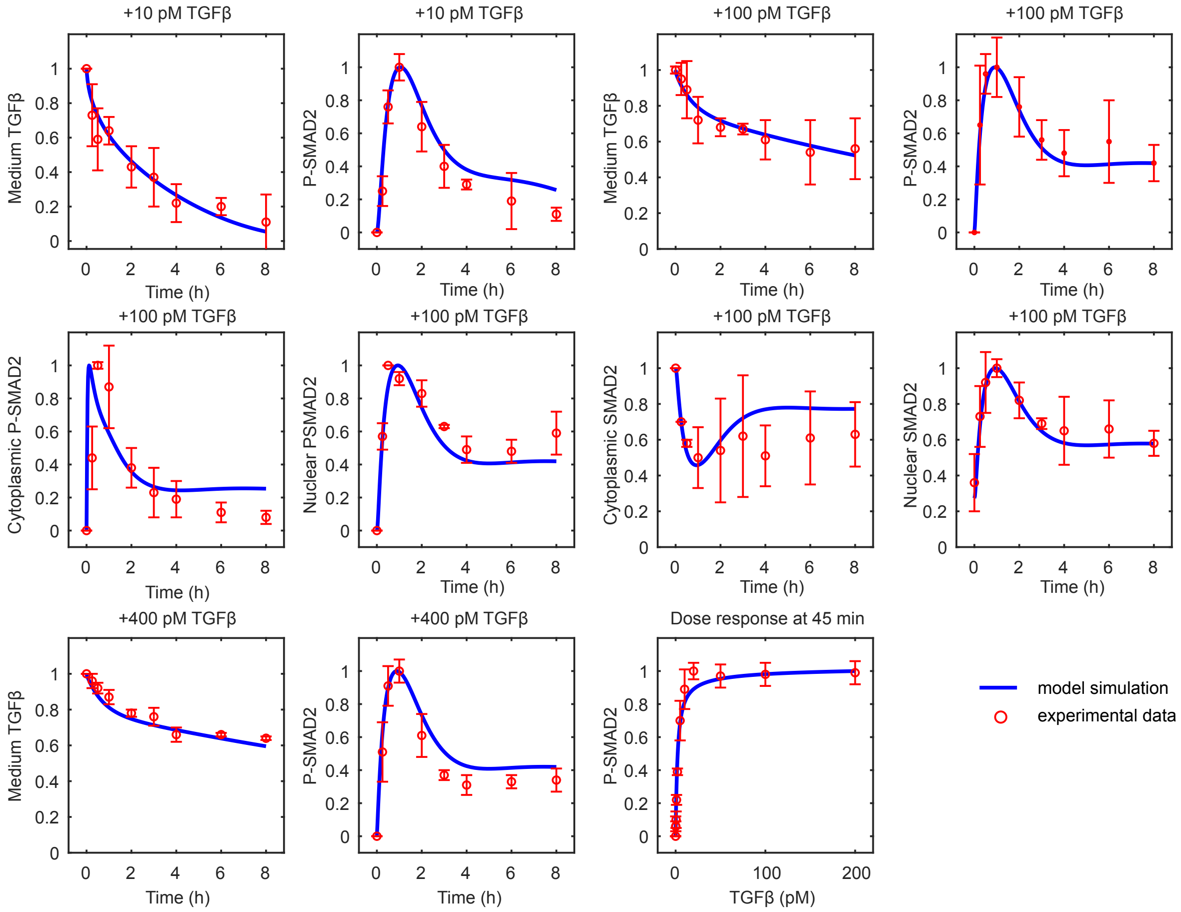

**Figure S4: The comparison of model simulation results and experimental data**

For comparison, the model simulation results and experimental data are normalized their maximum values in the corresponding data sets. The mean values and standard deviations of the normalized data from 3-5 experiments are shown.

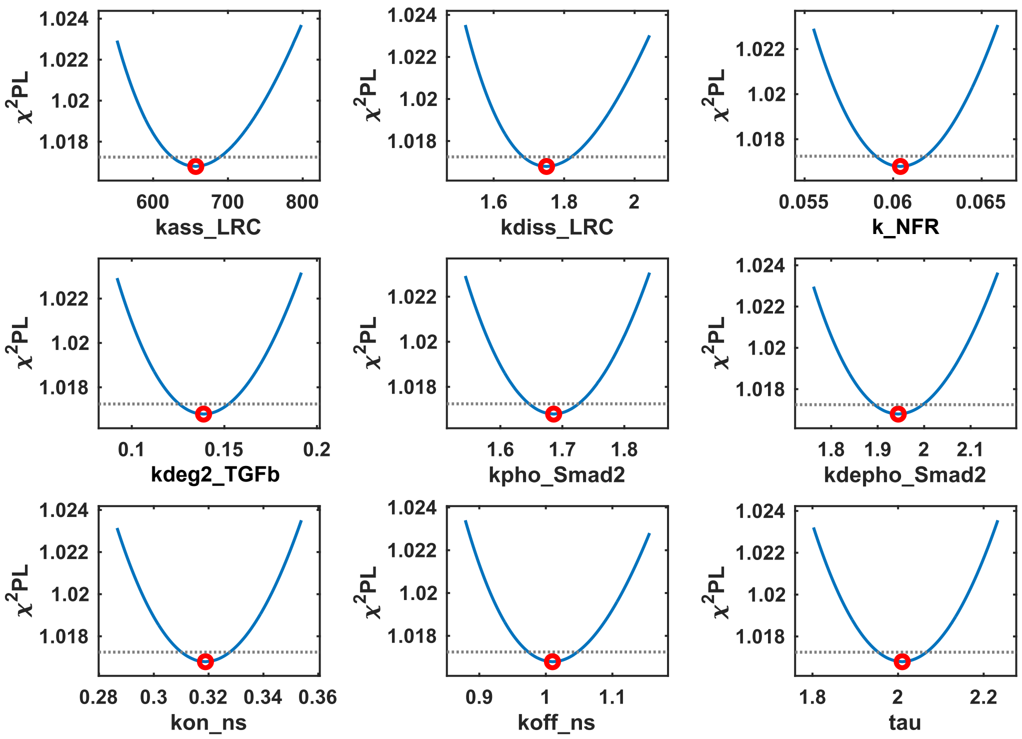

**Figure S5. Profile likelihood analysis of estimated parameter values**

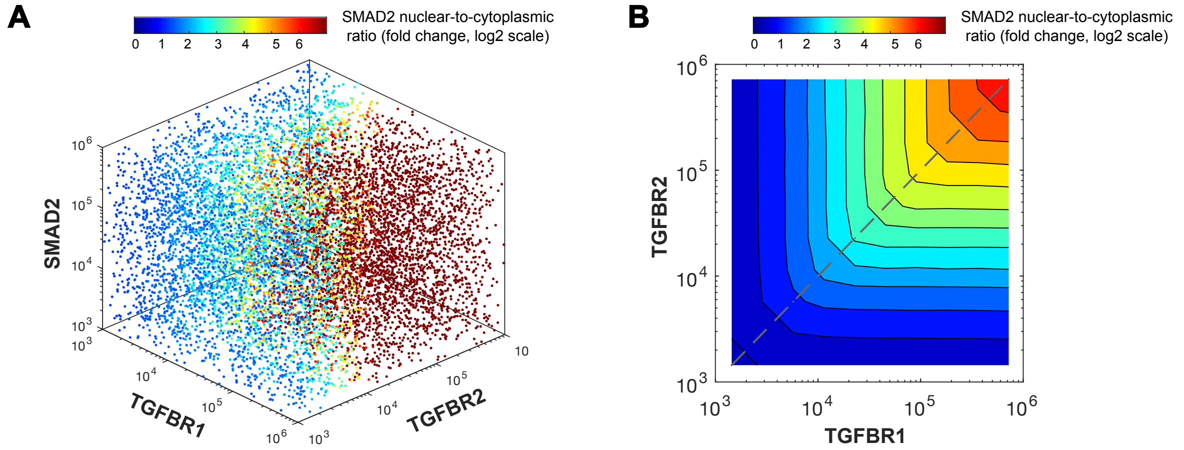

**Figure S6: Computational modeling of the ratio of nuclear-to-cytoplasmic SMAD2 in the variation space of TGF-β receptors and SMAD2.**

(A) Model predictions for the fold change in the ratio of nuclear-to-cytoplasmic SMAD2 (N2C) in the space of different amounts of TGFBR1, TGFBR2, and SMAD2 . Each point indicates a random set of protein amounts in the space. Colors indicate the fold change in the SMAD2 N2C ratio.

(B) Contour landscape of the fold change in SMAD2 N2C according to combinations of different amounts of TGFBR1 and TGFBR2. The dashed line indicates relationship between the fold change in SMAD2 N2C and the receptors when equal amounts of each receptor are present.

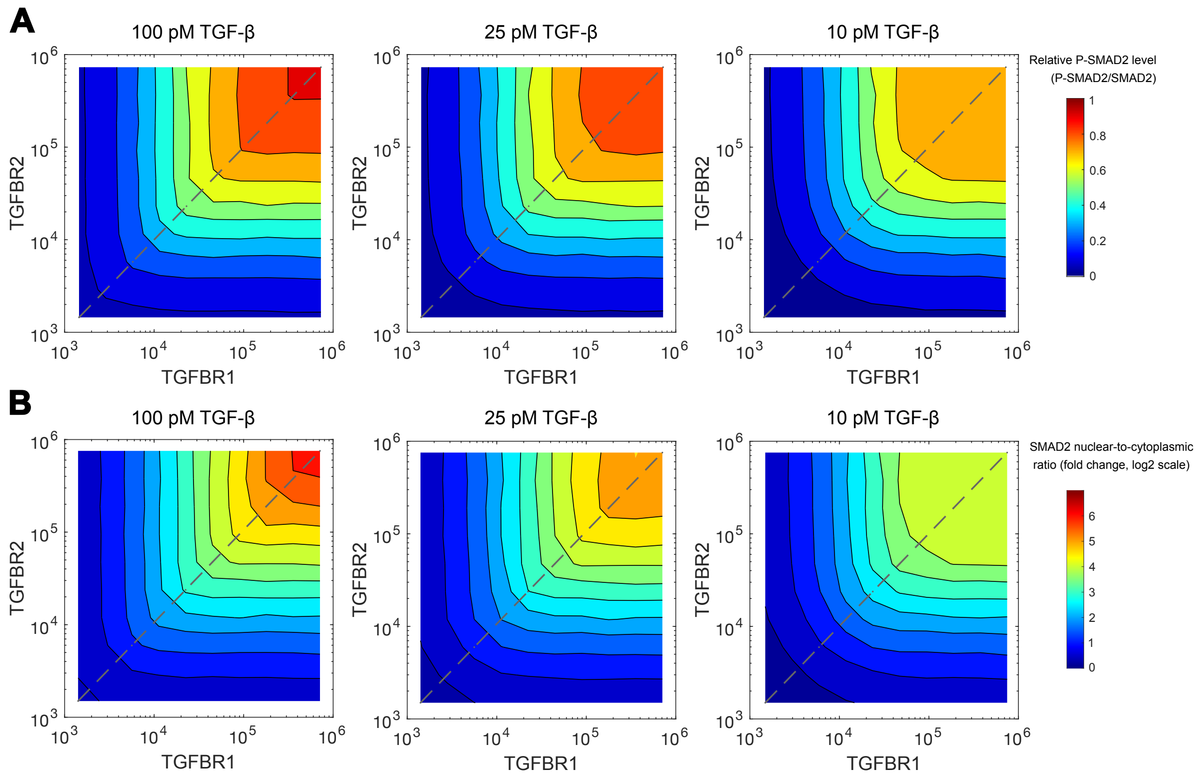

**Figure S7: Contour landscape of SMAD2 signaling responses in the variation space of TGF-β receptors and SMAD2 under different doses of TGF-β stimulations.**

(A) Contour landscape depicting the relative P-SMAD2 level based on combinations of different amounts of TGFBR1 and TGFBR2 with 100 pM, 25 pM, and 10 pM TGF-β stimulations.

(B) Contour landscape showing the fold change in SMAD2 N2C based on combinations of different amounts of TGFBR1 and TGFBR2 with 100 pM, 25 pM, and 10 pM TGF-β stimulation.

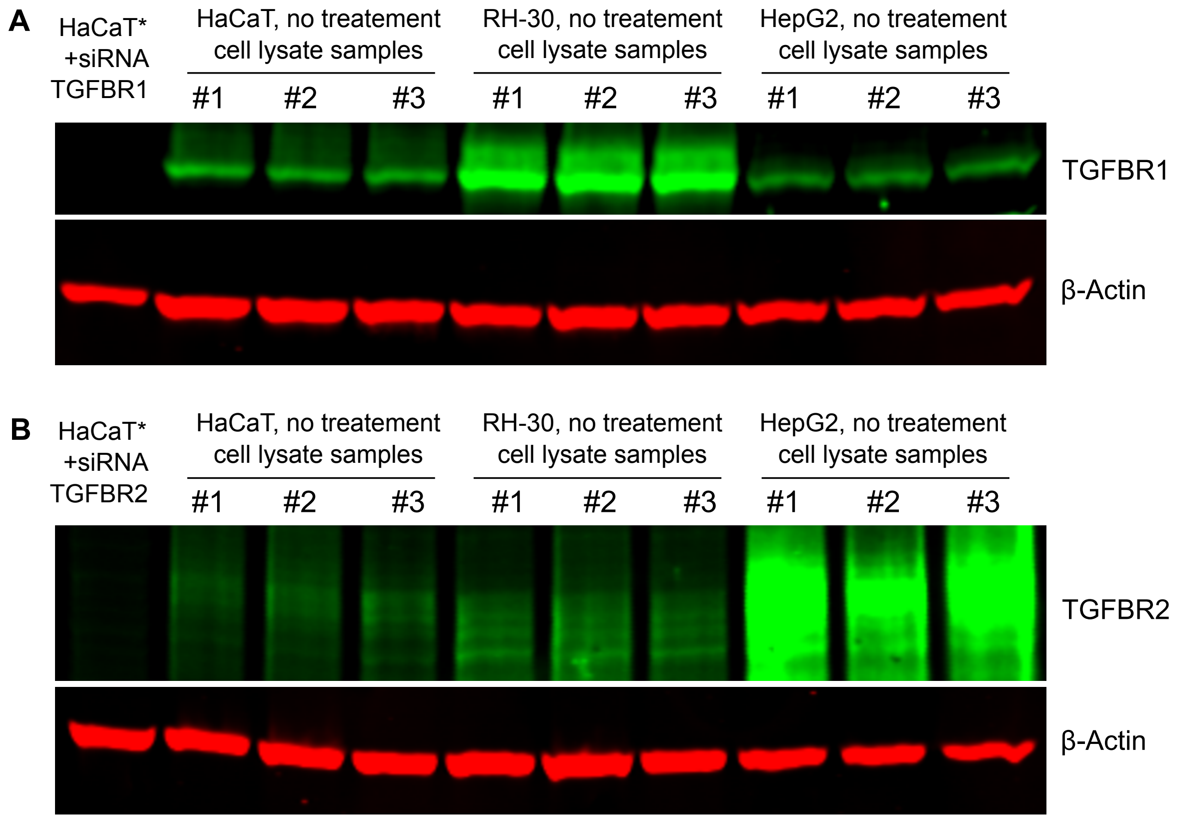

**Figure S8. TGFBR1 and TGFBR2 protein expression profiles in HaCaT, RH-30 and HepG2 cells**

Western blot for measuring relative abundance of TGFBR1 (A) and TGFBR2 (B) in HaCaT, RH-30 and HepG2 cells. The lysate samples from cells (#1, #2, #3) under normal growth conditions (no treatment) were loaded for each cell line. The lysate from HaCaT cells transfected with 100 nM TGFBR1 or TGFBR2 siRNA was loaded as a negative control (the first lane on the left) for the validation of the corresponding antibody.

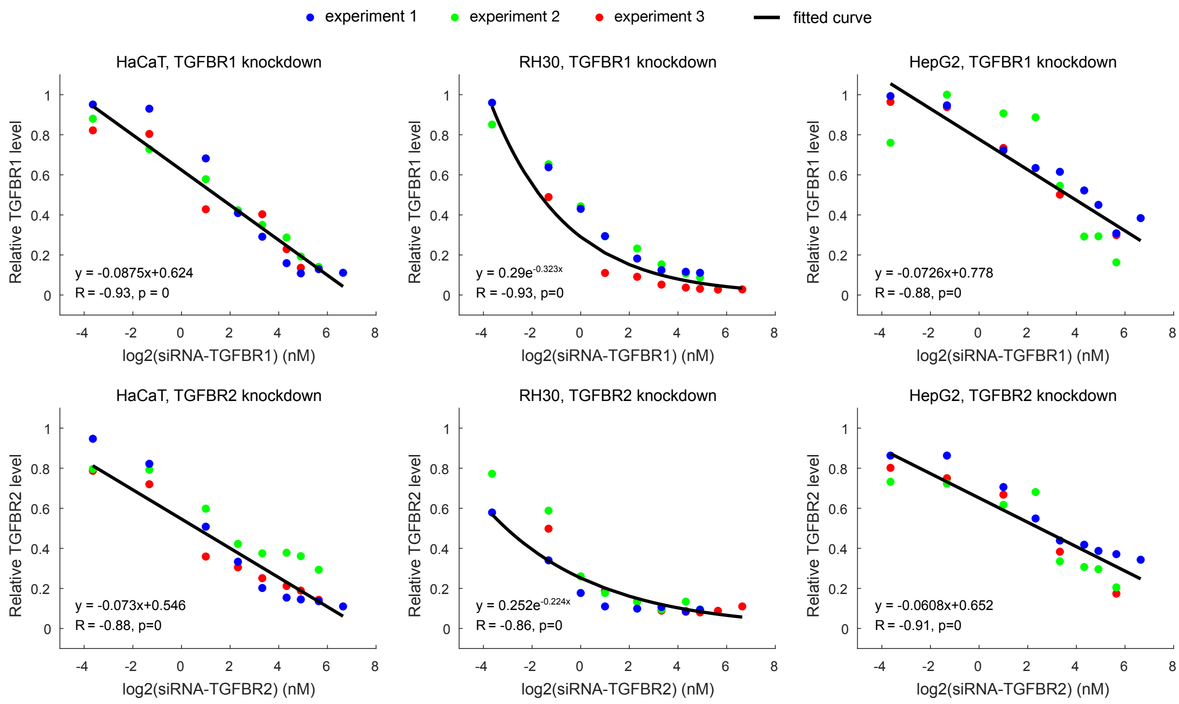

**Figure S9. Knockdown effect of TGFBR1 and TGFBR2 proteins using different concentrations of siRNAs in HaCaT, RH-30 and HepG2 cells**

The relative levels of TGFBR1 and TGFBR2 are normalized to the corresponding TGF-β receptor level of the non-targeting control (ntc) siRNA sample from the same experiment. The siRNA concentrations for TGFBR1 and TGFBR2 are plotted on a log2 scale, and the corresponding relative levels of TGFBR1 or TGFBR2 are displayed. The correlations (R-values and p-values) in the plots were quantified with Pearson correlation coefficients.

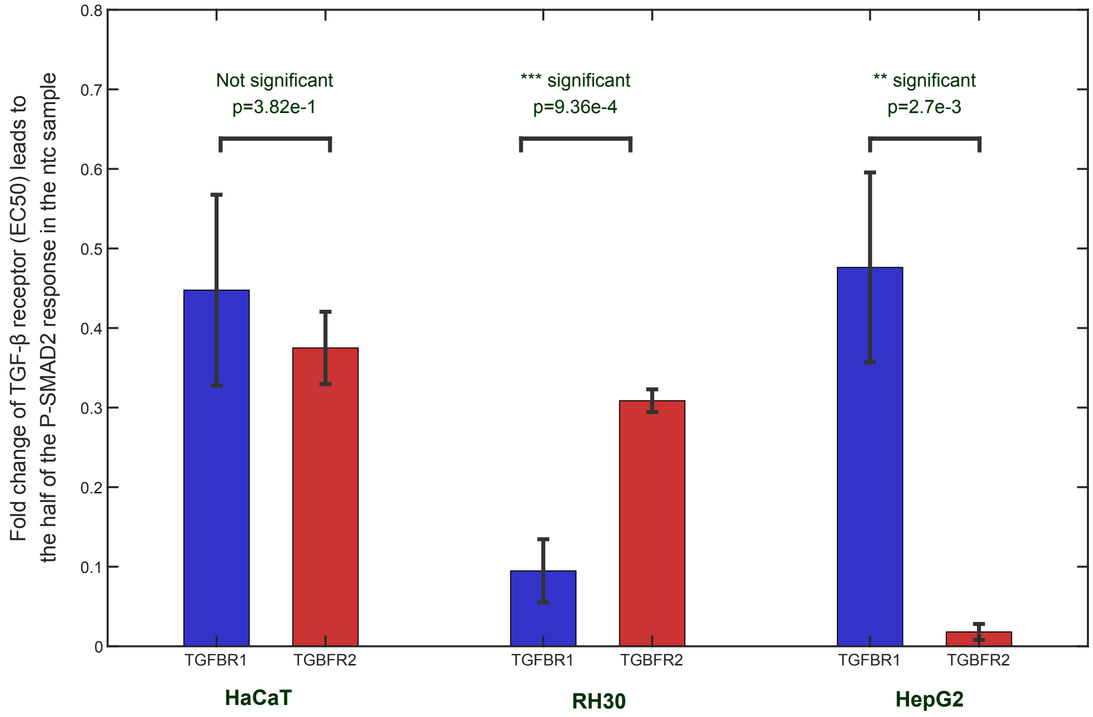

**Figure S10. Statistical analysis of TGF-β receptor fold-change effects leading to a 50% reduction in the P-Smad2 response compared to that in the non-targeting siRNA control group (EC50) during siRNA knockdown experiments.**

The EC50 values were determined by fitting Hill functions to the data shown in Figure 4. Presented here are the average EC50 values along with standard deviations, calculated from three independent experiments. Significance testing was conducted using a two-sample t-test, and significance levels are denoted as follows: ***$p < 0.001$, **$p < 0.01$.

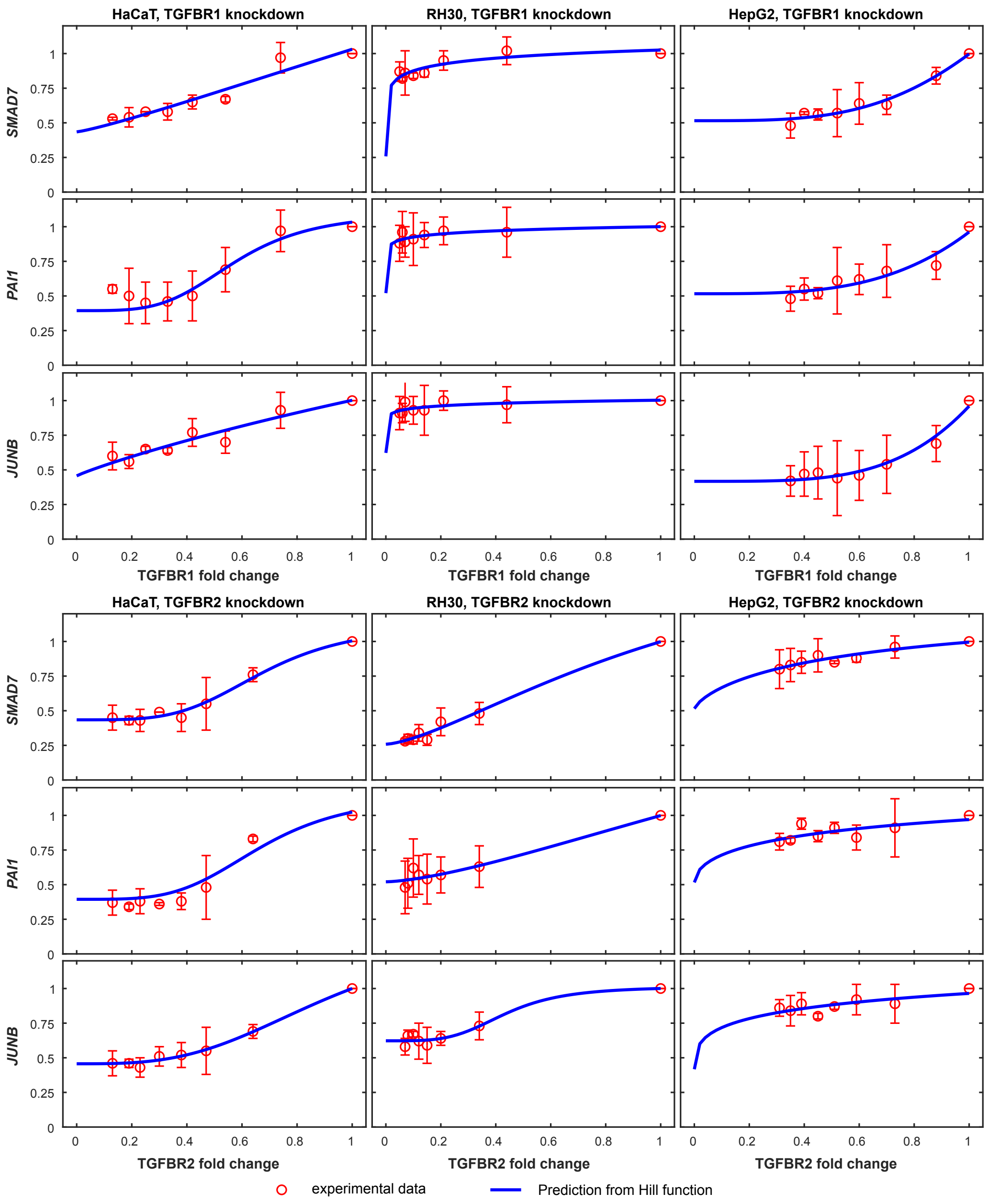

**Figure S11. Expression of TGF-β targeted genes in responses to the knockdown of TGF-β receptors in HaCaT, RH30 and HepG2 cells**

Different levels of each type of TGF-β receptor were achieved by knocking down with various concentrations of siRNA (non-targeting control, 0.4, 2, 5, 10, 20, 30, and 50 nM). The receptor levels were determined using calibration functions shown in Figure S9. The relative levels of TGF-β targeted genes (*SMAD7*, *PAI1*, and *JUNB*) were measured at 1 hour after 100 pM TGF-β stimulation and normalized to the non-targeting control (ntc) siRNA sample in the same experiment. The error bars represent the standard deviations of the quantified real-time PCR data from three biological replicates.

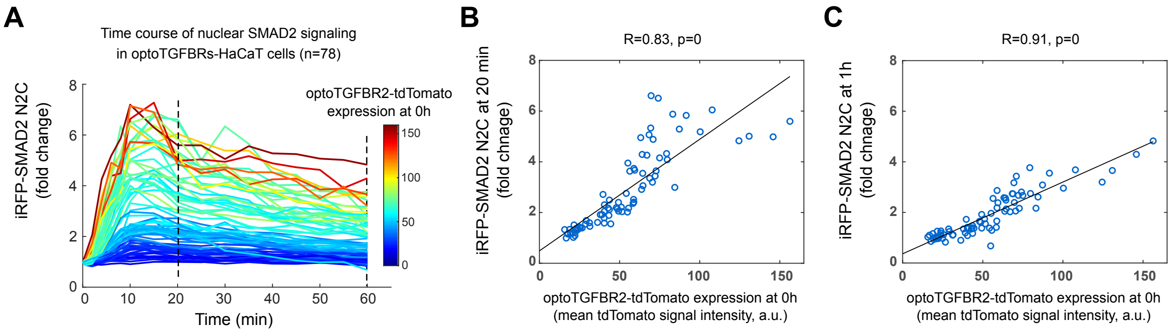

**Figure S12. Quantification of SMAD2 N2C fold change responses in HaCaT cells expressing similar levels of optoTGFBR1 and optoTGFBR2**

(A) Time-course profiles of iRFP-SMAD2 N2C fold change responses in individual optoTGFBRs-HaCaT cells after blue light stimulation.

(B) The plot of iRFP-SMAD2 N2C fold change responses at 20 min versus the amount of optoTGFBR2-tdTomato before blue light stimulation (0 h) in optoTGFBRs-HaCaT cells.

(C) The plot of iRFP-SMAD2 N2C fold change responses at 1 h versus the amount of optoTGFBR2-tdTomato before blue light stimulation (0 h) in optoTGFBRs-HaCaT cells.

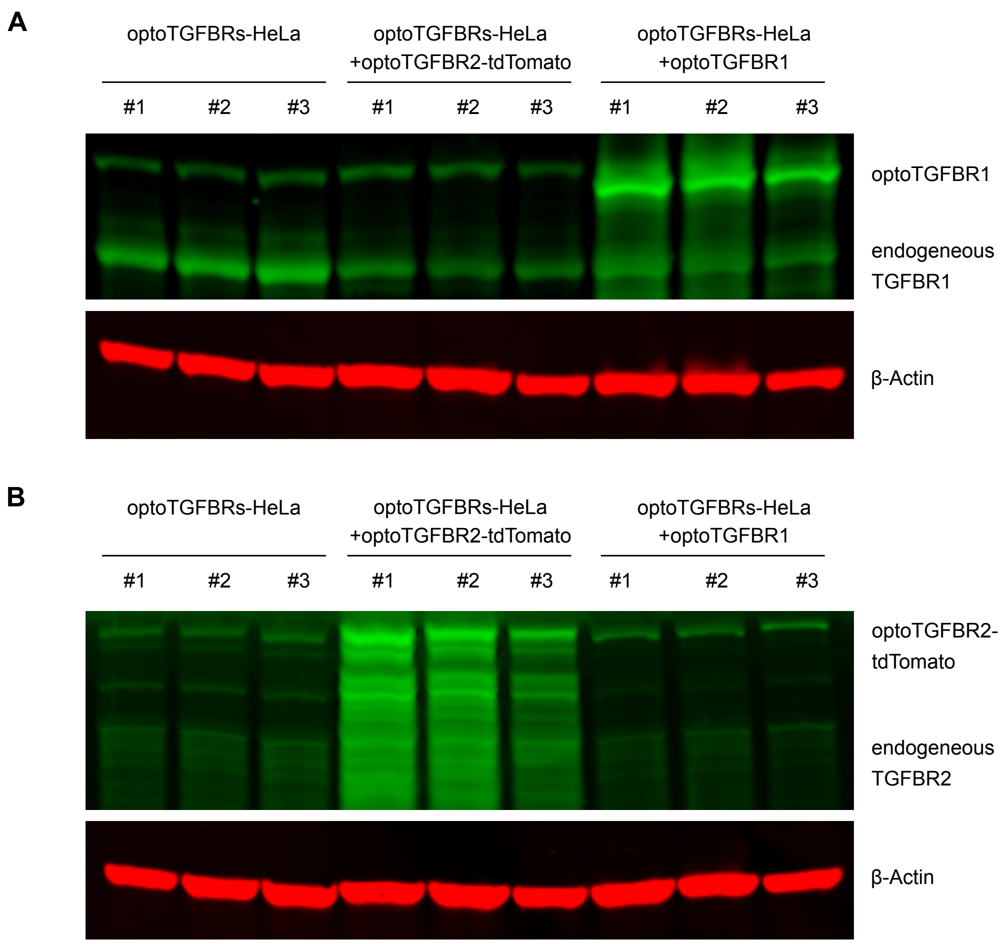

**Figure S13. Western blot analysis of optoTGFBR1 and optoTGFBR2 protein expression in optoTGFBRs-HeLa cells transfected with optoTGFBR1 or optoTGFBR2 plasmids**

Cell lysates were obtained from optoTGFBRs-HeLa cells, optoTGFBRs-HeLa cells transfected with optoTGFBR1 plasmids (optoTGFBRs-HeLa + optoTGFBR1), and optoTGFBRs-HeLa cells transfected with optoTGFBR2-tdTomato plasmids (optoTGFBRs-HeLa + optoTGFBR2-tdTomato). Immunoblotting was performed using antibodies for TGFBR1 (A) and TGFBR2 (B).
